# Young and old adult brains experience opposite effects of acute sleep restriction on the functional connectivity network

**DOI:** 10.1101/2025.07.25.666859

**Authors:** Josh Neudorf, Leanne Rokos, Kelly Shen, Brianne Kent, Anthony R. McIntosh

**Author notes:** Corresponding Author: Anthony R. McIntosh.

## Abstract

Chronic, long-term sleep loss is detrimental to brain health and cognitive ability. However, older adults are affected differently by acute, short-term loss of sleep than young and middle-aged adults. Older adults are more resilient to the effects of acute sleep loss and, depending on the cognitive domain, may be completely unaffected while younger adults suffer. To elucidate the brain network responses to sleep loss underlying these cognitive differences between age groups, we investigated the static and dynamic functional connectivity effects of sleep restriction (sleep limited to 3 hours) and how these effects differ between younger adults (20-30 years) and older adults (65-75 years). We found a functional connectivity subnetwork that was primarily strengthened in younger adults after sleep restriction but weakened in older adults after sleep restriction. Similar crossover interactions were consistently observed in further analyses of functional connectivity degree, modularity, and dynamic functional connectivity state fractional occupancy. Our findings demonstrate that the effect of sleep restriction on older adults is fundamentally different from younger adults. These results most strongly support the compensation theory of aging, which predicts a fundamental shift in the effects of sleep loss, rather than a mere dampening of the sleep benefits experienced by younger adults.

---

In fast-paced, work-centric cultures, it can sometimes feel as though there is not enough time in the day. We may resent our body’s daily need for sleep. In many cases, the combined demands of life, including work and family, lead to sacrificing sleep for the sake of finding more time. However, this loss of sleep is indeed a sacrifice, which can come at a cost. Beyond the well-known short-term effects of irritability and diminished cognitive ability, cumulative loss of sleep over the lifespan takes its toll, negatively impacting the brain and body. Poor sleep has the potential to increase the risk of cardiovascular disease, hypertension, diabetes, weight problems, colorectal cancer, and all-cause mortality, to name a few of the possible repercussions (see Medic et al., 2017 for a review). Poor sleep also adds up over time to negatively impact the brain. Middle-aged adults with short nights of sleep have more β-amyloid in their brains (Spira et al., 2013), which in many cases precedes cognitive decline and Alzheimer’s disease (Bateman et al., 2012). Indeed, many longitudinal studies have identified poor sleep in middle-age as a predictor of cognitive decline and Alzheimer’s disease later in life (Ferrie et al., 2011; Kulmala et al., 2013; Lobo et al., 2008; Stenfors et al., 2013; Virta et al., 2013).

The acute, short-term effects of sleep loss on cognitive performance and the brain are more nuanced than the chronic, long-term effects. Young and middle-aged adults’ attention, executive control, working memory, episodic memory, and mood are impacted negatively when sleep is missed (e.g., Boland et al., 2017; Krieg et al., 2001; Kronholm, 2012; Pilcher & Huffcutt, 1996; Regestein et al., 2004; Scullin & Bliwise, 2015; Sternberg et al., 2013). However, there are also temporary positive effects of skipping or limiting a night of sleep. For example, across multiple studies, the antidepressant effects of acute sleep deprivation have been well replicated. Around half of people with depression symptoms respond positively to acute sleep deprivation, noting a reduction in their symptoms (see Boland et al., 2017 for a review). Furthermore, there are many effects of acute sleep deprivation that have been observed in younger adults but have not been replicated in older adults or are markedly weaker. Across multiple cognitive domains and many studies, it has been demonstrated that older adults are resilient to the cognitive effects of sleep deprivation (Gerhardsson et al., 2019; Scullin & Bliwise, 2015; Sutter et al., 2012). Older adults are also more resilient to the negative effects of sleep deprivation on mood (healthy adults generally respond with more negative mood in response to sleep deprivation, as opposed to the exception noted above for some people with depression symptoms; Schwarz et al., 2019).

There have been a number of theories developed to explain why older adults are more resilient to sleep loss than younger adults (see Scullin & Bliwise, 2015 for a review). The most prominent of these theories include the reduced sleep need, functional weakening, and compensation theories. The reduced sleep need theory suggests that older adults need less sleep than they did as younger adults. Whereas earlier in the lifespan sleep was an important process for maintaining cognitive ability, as the amount of learning decreases with age, so too does the need for sleep to consolidate that learning (Cirelli, 2012; Feinberg, 2000). A second theory, the functional weakening theory, suggests that the aging brain undergoes multiple negative changes that undermine the ability of sleep to perform the restorative, constructive role it once held (Edinger et al., 2000; Kronholm, 2012; McCrae et al., 2012; Pace-Schott & Spencer, 2011; Schmidt et al., 2012; Scullin, 2013; Spiegel et al., 1986). For this reason, not only do older adults not need additional sleep to maintain their baseline cognitive performance, but an increase in sleep will not produce benefits because the brain architecture that once supported the sleep-cognition relationship has deteriorated. Finally, the compensation theory suggests the brain undergoes fundamental adaptations with age to compensate for the changing roles of particular sleep stages for cognition, which affect resilience to sleep loss (Nemeth et al., 2013; Park & Reuter-Lorenz, 2009; Peters et al., 2008; Salas et al., 2014; Spencer & Pace-Schott, 2013). This theory predicts that the effects of sleep loss will be fundamentally different for older adults, rather than simply being a less pronounced version of what is observed for younger adults. Going beyond the cognitive effects of sleep loss across the lifespan to investigate the brain changes that underlie these differences may provide additional insight into which of these theories are most consistent with the evidence.

Researchers have studied young adult brains before and after loss of sleep, producing several interesting findings. During verbal tasks, the brain compensates for sleep deprivation by increasing frontal lobe activation, while frontal lobe activation is decreased in the absence of task demands (Drummond et al., 2000). This effect on the frontal lobe appears to exhibit inter-individual differences, whereby fatigue-resistant individuals who do not suffer as much of a loss in cognitive performance from sleep loss showed greater activation in the frontal lobe during a working memory task than fatigue-vulnerable individuals (Caldwell et al., 2004). Furthermore, individuals with strong connectivity between the hippocampus and thalamus after sleep restriction also have preserved short-term memory task performance (Chengyang et al., 2017). A brain network analysis of sleep deprivation has revealed an increase in the diversity of connections to connector hub regions after sleep deprivation, associated with increased global efficiency and acting as a potential compensatory reaction (Tian et al., 2024). Many other differences have been observed between normal and deprived sleep conditions, measured using functional stability (Huang et al., 2022), connectivity between large-scale brain networks (Chee & Zhou, 2019; Dai et al., 2020; Yeo et al., 2015), whole-brain modularity (Ben Simon et al., 2017), and dynamic functional connectivity measures of state dwell time and transition probability (Xu et al., 2018). Together, these findings in younger adults suggest that sleep deprivation impacts an individual’s brain network in many ways, but little is known about how the effects of sleep deprivation present in the brain networks of older adults. This research has primarily been performed on young adult brains, and comparing these effects across age groups remains a gap in the literature. One study comparing the effects across age groups focused on the interaction between sleep restriction (sleep limited to 3 hours) and age on the functional connectivity between a small number of large functional networks but did not identify a significant interaction (Nilsonne et al., 2017). However, there are more advanced methods that could be used to investigate the brain as a finer-scaled network with hundreds of regions to identify network changes at the scale of the regions and connections between regions and at the whole-brain scale utilizing a modularity analysis. Furthermore, dynamic functional connectivity changes have not been investigated in this context, which would provide further insights.

Functional connectivity analyses investigate the coactivation of regional brain activation, using, for example, functional magnetic resonance imaging (fMRI) blood oxygen level-dependent (BOLD) signal, to calculate a measure of functional correlation between brain regions. Graph theory is the field of mathematics focused on networks and has been applied to brain network data with tremendous success (Sporns, 2018). Graph-based analysis methods have also been developed expressly for the comparison of population groups’ brain network data, including the network-based statistic for comparing parcellated brain connectivity matrices (Zalesky et al., 2010) and multivariate distance matrix regression for comparing voxelwise brain connectivity matrices (Shehzad et al., 2014). In addition to studying static functional connectivity as described above, the dynamics of functional connectivity should also be investigated, as it has been demonstrated that functional connectivity patterns rapidly change over time, even at rest, and that these dynamics are meaningful as they are associated with cognitive ability (Cabral et al., 2017; Cohen, 2018; Hutchison et al., 2013; Neudorf et al., 2025) and disease (Cohen, 2018; Dautricourt et al., 2022; Jin et al., 2023; Zang et al., 2024). We propose to use these methods to investigate how the functional brain network effects of acute sleep restriction differ between younger and older adults. We hypothesize that there should be a significant interaction between the effects of acute sleep restriction and age, considering the consistent findings that older adults are less affected cognitively by acute sleep deprivation and restriction than younger adults. The theories that have been developed to explain this phenomenon produce different hypotheses for the nature of this interaction. The reduced sleep need and functional weakening theories propose reduced beneficial effects of sleep and would predict a reduced brain network effect of sleep restriction in older adults. On the other hand, the compensation theory proposes that older adults undergo a fundamental change in how the brain relies on the various sleep stages in order to support cognitive abilities, such as memory consolidation. Under the assumptions of this theory, we expect to see more of a fundamental shift in the effect of sleep restriction on the brain network rather than simply a dampening of the effect seen in younger adults. A shift of this kind would not be surprising, given the fundamental differences from the young adult brain that we have observed in the functional communication and network organization of the aging brain (Heisz et al., 2015; Neudorf et al., 2024, 2025).

## Methods

### Participants

Details about recruitment, ethics, and the full dataset can be obtained from the original dataset paper and a follow-up analysis paper (Nilsonne et al., 2016, 2017). Data were downloaded from OpenfMRI (https://openfmri.org/dataset/ds000201/). Participants were recruited in two different age groups that were either between the ages of 20-30 (young adults; YA) or between the ages of 65-75 (old adults; OA). The participants engaged in two separate scanning sessions on separate days, in which they either had a full night’s sleep or sleep restricted to 3 hours. The order of these sessions was counterbalanced, and took place approximately 1 month apart. Among other screening criteria, these participants had no past psychiatric or neurological illness and did not exhibit insomnia symptoms or excessive snoring.

### MRI Acquisition

Full details about MRI acquisition can be obtained from the original paper (Nilsonne et al., 2017). A General Electric Discovery 3T MRI was used. For the resting state fMRI (rs-fMRI), echo-planar images were acquired with a flip angle of 75 degrees, TE of 30 ms, TR of 2.5 s, field of view of 28.8 cm, and slice thickness of 3 mm, acquiring 49 slices that were interleaved from the bottom up. In our analysis, from each participant 173 volumes (totaling 7 minutes and 12.5 seconds) were used in order to equate the total number of volumes. The T1-weighted structural images utilized a sagittal BRAVO sequence with field of view of 24 cm and slice thickness of 1 mm acquired from the bottom up.

### Preprocessing

Preprocessing of the MRI data utilized TheVirtualBrain-UK Biobank pipeline (Frazier-Logue et al., 2022), based on the UK Biobank pipeline (Littlejohns et al., 2020), and using the FMRIB Software Library (FSL; Jenkinson et al., 2012). TheVirtualBrain-UK Biobank pipeline accounts for issues related to atrophy in aging brains using quality control methods outlined by Lutkenhoff et al. (2014). In addition, the pipeline produces parcellation-based time-series data from the fMRI volumes which were used in the functional connectivity analyses. The data were parcellated using a combined atlas of the Schaefer 200 region atlas (Schaefer et al., 2018) and the subcortical Tian atlas (Tian et al., 2020). The subcortical regions included regions from the Tian Scale 1 atlas but with the hippocampus represented by the Tian Scale 3 atlas definition, with the two head divisions combined into a single parcel. A custom-trained *FSL FIX* classifier was provided to the pipeline, trained on 10 scans from the first scanning session and 10 scans from the second scanning session, balancing the number of YA and OA participants and male and female participants, with *FIX* set to “aggressive” artefact removal (Griffanti et al., 2014).

The neuroimaging outputs were also subjected to manual quality control, which included manual inspection of brain extraction, segmentation, registration, masks, fMRI time-series plots, and FC matrices. Structural T1 images were checked for motion artefacts and poor brain extraction, registration was checked for proper alignment, fMRI time-series were inspected for residual motion artifacts, and FC matrices were inspected for poor separation of the values into intra- and inter-hemispheric quadrants and prominent “banding”, whereby a single region exhibits fairly constant values indicating motion artifacts and/or poor registration. Following the previous publication with this data, participants passing the previous quality control steps were checked to ensure that no more than 25% of volumes had framewise displacement greater than 0.5 mm. There were 48 YA and 36 OA in the sample, and following our quality control of the magnetic resonance imaging (MRI) data, we included 23 YA (11 female) and 19 OA (9 female).

Time series data from the pipeline was then bandpass filtered to remove frequencies outside of the range between 0.01 Hz and 0.1 Hz. To calculate the FC matrices, the time series were Z-scored across time separately for each region, the Pearson’s R correlation was calculated between each pair of regions, and then a Fisher Z-transformation was applied.

### Dynamic Functional Connectivity

The rs-fMRI data was analyzed using a dynamic functional connectivity (dFC) analysis approach called Leading Eigenvector Dynamics Analysis (LEiDA; Cabral et al., 2017). This method computes an FC matrix at each time point of the rs-fMRI scan using phase coherence connectivity (e.g., Deco et al., 2017; Deco & Kringelbach, 2016; Glerean et al., 2012; Ponce-Alvarez et al., 2015). In contrast to other dFC methods that compare the full FC matrices across timepoints, LEiDA first computes the leading eigenvector of each FC matrix, making the method less susceptible to noise and better able to detect the recurrence of a particular state. The dFC matrices are separated into distinct states by applying a clustering analysis on the leading eigenvectors for all subjects and timepoints, resulting in states that are common to all subjects.

The ideal number of states was chosen based on an evaluation of the clustering analysis that maximized the Dunn’s score (Dunn, 1973), average Sihouette coefficient (Rousseeuw, 1987), and Calinski–Harabasz index (Caliński & Harabasz, 1974). With these states defined, each timepoint was then labelled according to which state the participant’s brain network was in at that timepoint, which allowed for calculation of the fractional occupancy (FO) of each state (probability of that state occurring at any given time) and the transition probability matrices (probability of the brain state changing from a specific state to another, or maintaining the same state, represented as a K×K matrix where K is the total number of states).

### Modularity

Modularity indicates the degree to which a network can be separated into distinct modules (i.e., communities). High modularity indicates that the network has strong connectivity within modules but weaker connectivity between modules. This measure was calculated using the Brain Connectivity Toolbox (BCT; Rubinov & Sporns, 2010) function *community_louvain*, with asymmetric treatment of negative weights from the FC matrices. To test that the detected communities were more modular than would be expected by chance, 1000 null networks were produced using the *null_model_und_sign* function, which preserves weight and degree distributions as well as the approximated strength distributions. All networks’ modularity values surpassed every permuted null value.

### Signal Variability

Grey matter signal variability was calculated by calculating the standard deviation of each of the grey matter voxelwise bandpass filtered (0.01 to 0.1 Hz) signals, then calculating the mean across all voxels. This value was then log-transformed to address skewness.

### Partial Least Squares (PLS)

Multivariate partial least squares (PLS) analysis (McIntosh & Lobaugh, 2004) was used to identify latent variables (LVs), each containing weights that describe the relationship of brain measures (e.g., FC, FC degree, fractional occupancy, or transition probability) with age (between-subjects) and sleep condition (within-subjects). Mean-centered PLS analyses were performed by categorizing scans into 4 groups based on their age (YA or OA) and sleep condition (Restricted or Normal). Additional models also included sex as a variable, resulting in a total of 8 groups of scans (11 Female YA Restricted Sleep, 11 Female YA Normal Sleep, 12 Male YA Restricted Sleep, 12 Male YA Normal Sleep, 9 Female OA Restricted Sleep, 9 Female OA Normal Sleep, 10 Male OA Restricted Sleep, and 10 Male OA Normal Sleep). PLS uses singular value decomposition to project the data matrix onto orthogonal LVs (similar to canonical correlation analysis). The significance of the identified LVs was determined via permutation testing. We report only the most reliable PLS weights as determined by bootstrap resampling, which is used to calculate bootstrap ratios (BSR), which are the ratios of the PLS weights (saliences) to their standard errors as determined by bootstrap resampling (Kovacevic et al., 2013; McIntosh & Lobaugh, 2004). The resampling procedures were performed using 1000 iterations for each. For the mean-centered PLS, brain scores are derived for each LV using the dot-product of the PLS weights for the brain metric and the values of the metric for each participant. The brain scores are similar to factor scores and can indicate the degree to which a subject or group shows the pattern captured in the LV. In the present paper, we use these to convey the relative difference in LV expression between groups, using the mean brain score and the bootstrap estimated standard error.

### Network-Based Statistic (NBS)

A network-based statistic (NBS; Zalesky et al., 2010) approach was used to identify an age group related subnetwork from the functional connectivity sleep difference network (values from normal sleep subtracted from those obtained during restricted sleep). Each connection in this sleep difference network was used to compute an independent samples t-test between the two groups, which was thresholded at *p* < .01 (*t* > 2.7). A significant subnetwork was then checked for based on a permutation test with 5000 permutations comparing the empirical subnetwork size (number of involved regions) to those observed with permuted group labels.

### Multivariate Distance Matrix Regression (MDMR)

Multivariate distance matrix regression (MDMR; Shehzad et al., 2014) was used to identify voxelwise differences in the FC connectivity profiles between age groups and sleep conditions. Voxels were resampled to 4 mm^3^ in MNI standard space and grey matter voxelwise FC matrices were computed in the same manner used for the parcellation-based FC matrices. Group comparisons between YA and OA were tested on the voxelwise FC sleep difference network, and sleep condition comparisons were tested on the voxelwise FC network for YA and OA separately, using 15,000 pseudo-F-statistic permutations and 500 cluster size permutations. Surface visualizations of these results were produced using the *neuromaps* (Markello et al., 2022) implementation of volume-to-surface registration as proposed by Buckner et al. (2011) and implemented by Wu et al. (2018).

## Results

### Functional Connectivity

A mean-centred PLS analysis was performed with FC as the dependent variable and independent variables of age group (YA vs. OA) and sleep condition (Restricted vs. Normal). This analysis identified one significant LV (*p* = .041) associated with a crossover interaction between age and sleep, whereby the effect of sleep restriction occurred in opposite directions for YA and OA (see Figure 1). As seen by the lines showing the sleep effect for individuals by sex, for the YA males, all but two of these individuals followed the pattern identified by the LV, while YA females were much more inconsistent. For OA, all but one of the males followed the pattern identified by the LV, and all of the females followed the pattern (see Figure 1). The functional connections reliably associated with this LV were primarily negatively associated (682/710 connections; 96.056%), indicating that sleep restriction in OA was associated with reduced connectivity among these regions but increased connectivity in YA (see Figure 2). A smaller number of connections were positively associated with the LV, indicating that these connections had increased connectivity during sleep restriction in OA but decreased connectivity during sleep restriction in YA. These connections were primarily made up of connections involving the occipital lobe. The most highly connected regions within this subnetwork of identified regions included the left hemisphere (LH) orbitofrontal cortex (OFC), superior parietal lobule (SPL), and caudate, and the right hemisphere (RH) dorsal prefrontal cortex (dPFC), somatosensory cortex, auditory cortex, and posterior thalamus. Of the 710 identified connections, 21.972% were within the LH, 23.662% were within the RH, and 54.366% were interhemispheric. Looking at the mean connectivity difference between restricted and normal sleep for these connections, OA exhibited a negative difference for all connections with negative BSRs and a positive difference for all connections with positive BSRs, consistent with the pattern identified by the LV. For YA, the majority (87.042%) of connections exhibited a positive difference for the negative BSRs and a negative difference for the positive BSRs, consistent with the pattern identified by the LV (see Figure 3).

**Figure 1.**
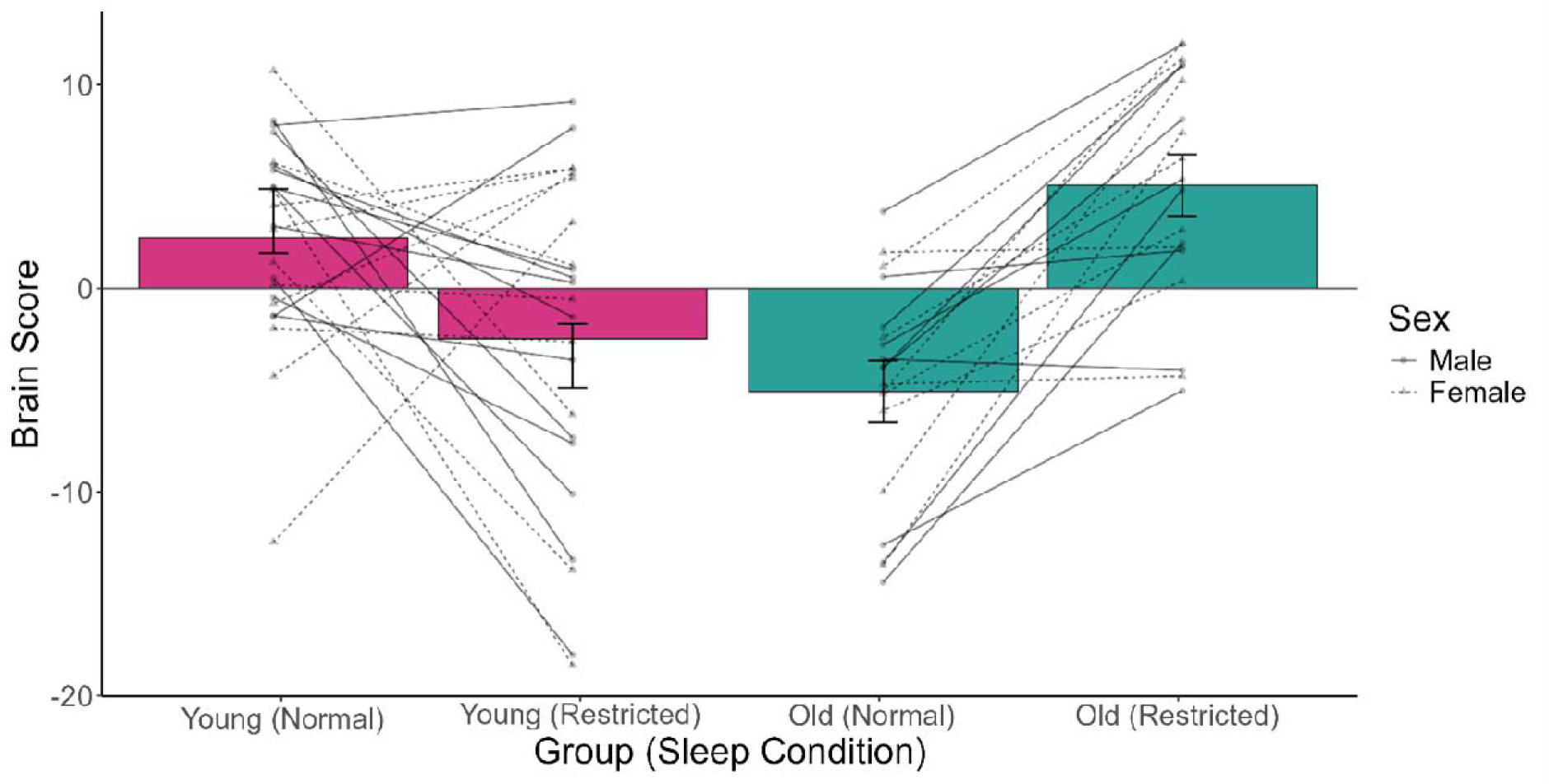
Mean brain scores from FC mean-centred PLS LV, for participants in the young and old age groups, in restricted and normal sleep conditions. Error bars represent 95% confidence intervals. Solid (male) and dashed (female) lines depict the change in the individual brain score values between sleep conditions.

**Figure 2.**
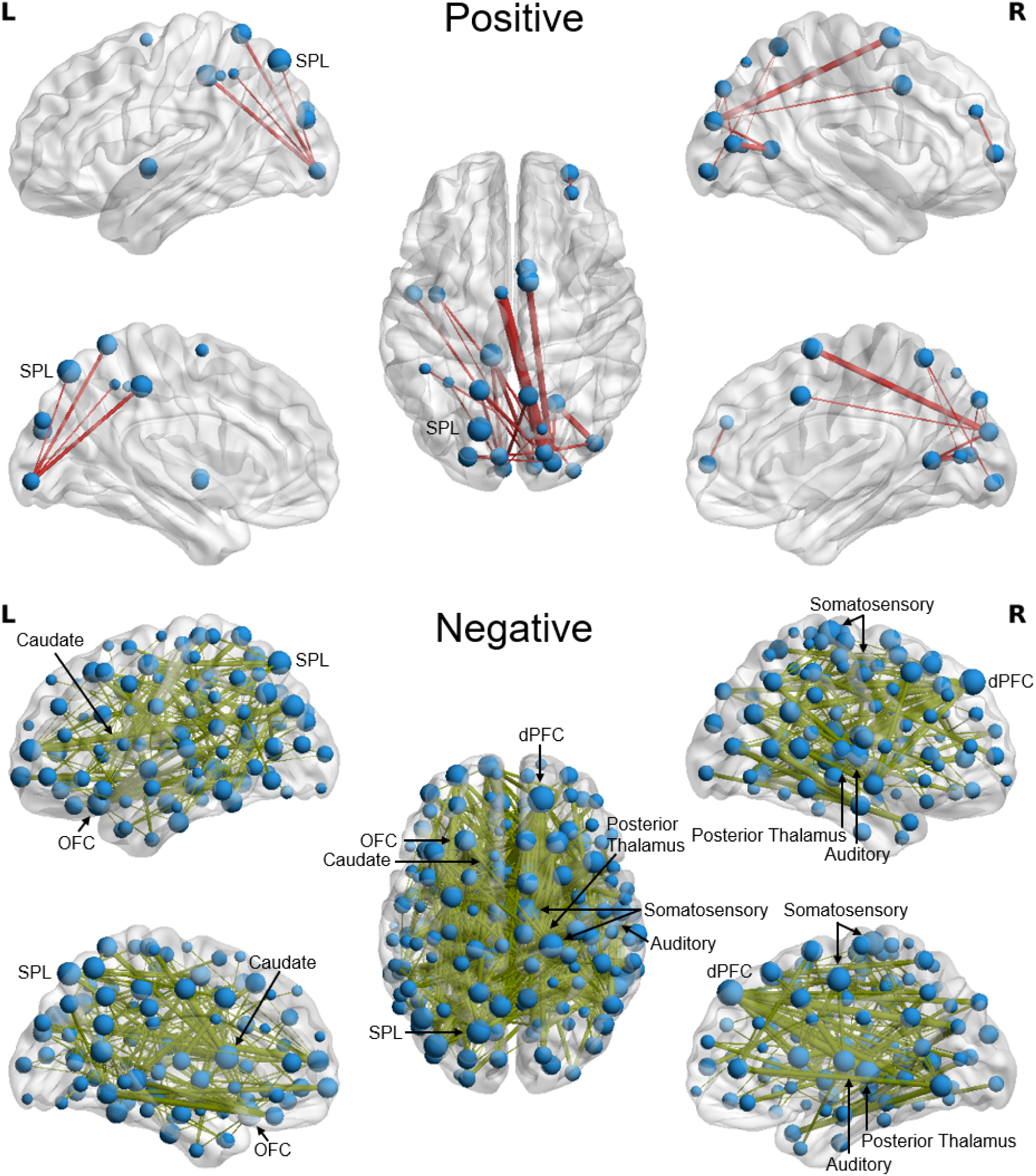
Reliable bootstrap ratios from FC mean-centred PLS LV with independent variables of age and sleep condition, depicted separately for positive and negative values. Connection size indicates the magnitude of the BSR. Highest degree regions are labelled (SPL: superior parietal lobule; OFC: orbitofrontal cortex; dPFC: dorsal prefrontal cortex).

**Figure 3.**
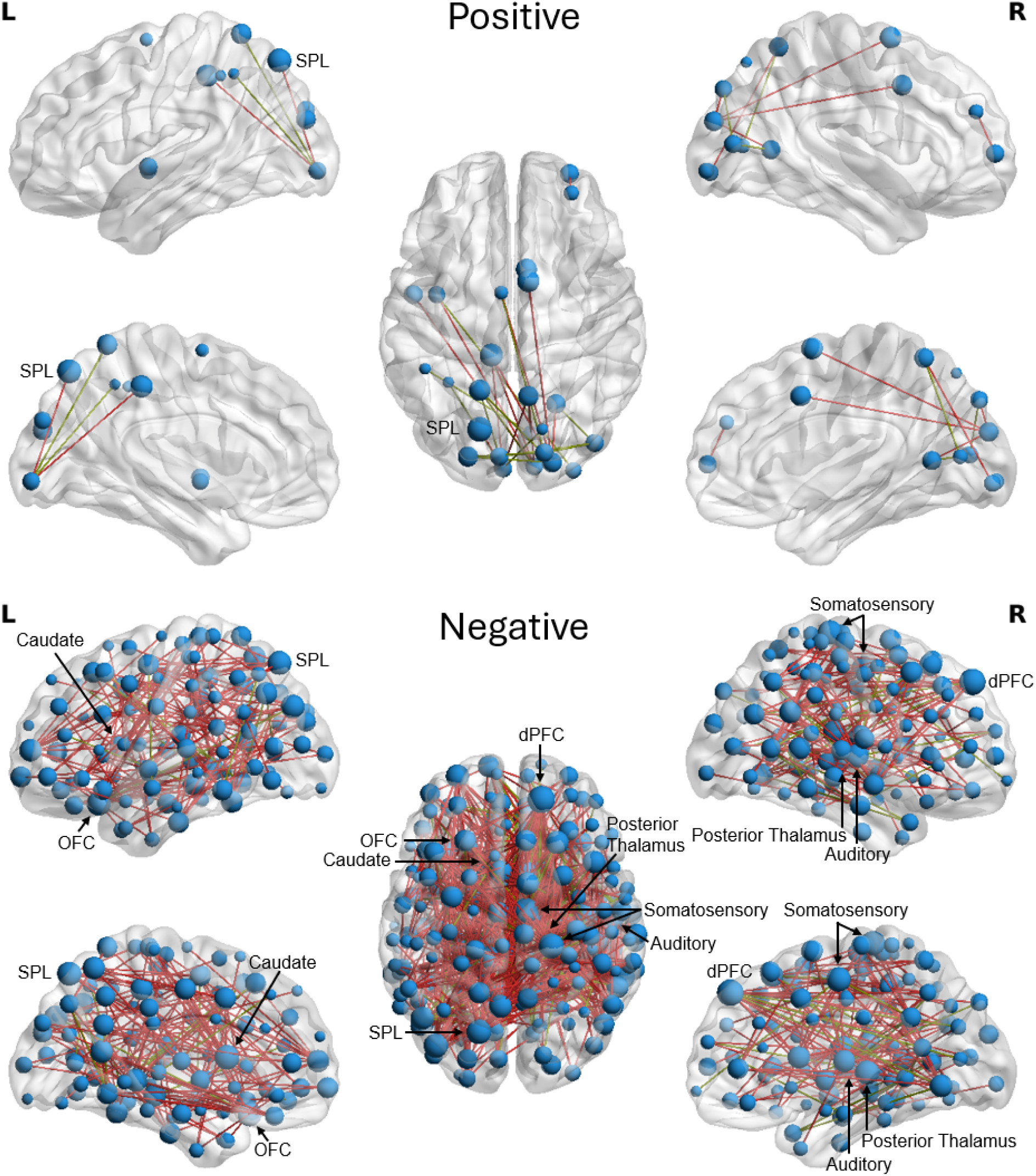
Mean connectivity difference between restricted and normal sleep conditions in YA for connections identified as reliable by the FC mean-centred PLS LV. Highest degree regions are labelled (SPL: superior parietal lobule; OFC: orbitofrontal cortex; dPFC: dorsal prefrontal cortex).

A similar analysis was conducted on the functional connectivity data using the network-based statistic (NBS; Zalesky et al., 2010). The functional connectivity sleep difference network was calculated by subtracting the normal sleep condition from the restricted sleep condition connectivity, and NBS was used to identify a subnetwork within this sleep difference network that differed between YA and OA groups (*p* = .040). The results of this analysis are shown in Figure 4, where positive values represent connections with a larger positive sleep effect for OA than YA and negative values represent connections with a higher magnitude negative sleep effect for OA than YA. This analysis identified a significant “difference of differences” (i.e., interaction) network similar to the network identified by the PLS. The most highly connected regions identified in this subnetwork included the LH OFC and RH dPFC, somatosensory cortex, auditory cortex, and posterior thalamus, which were highlighted in the PLS analysis, as well as the LH lateral ventral prefrontal cortex (lvPFC) and precuneus, and the RH globus pallidus. This subnetwork of 539 connections consisted of 20.965% LH connections, 22.820% RH connections, and 56.215% interhemispheric connections.

**Figure 4.**
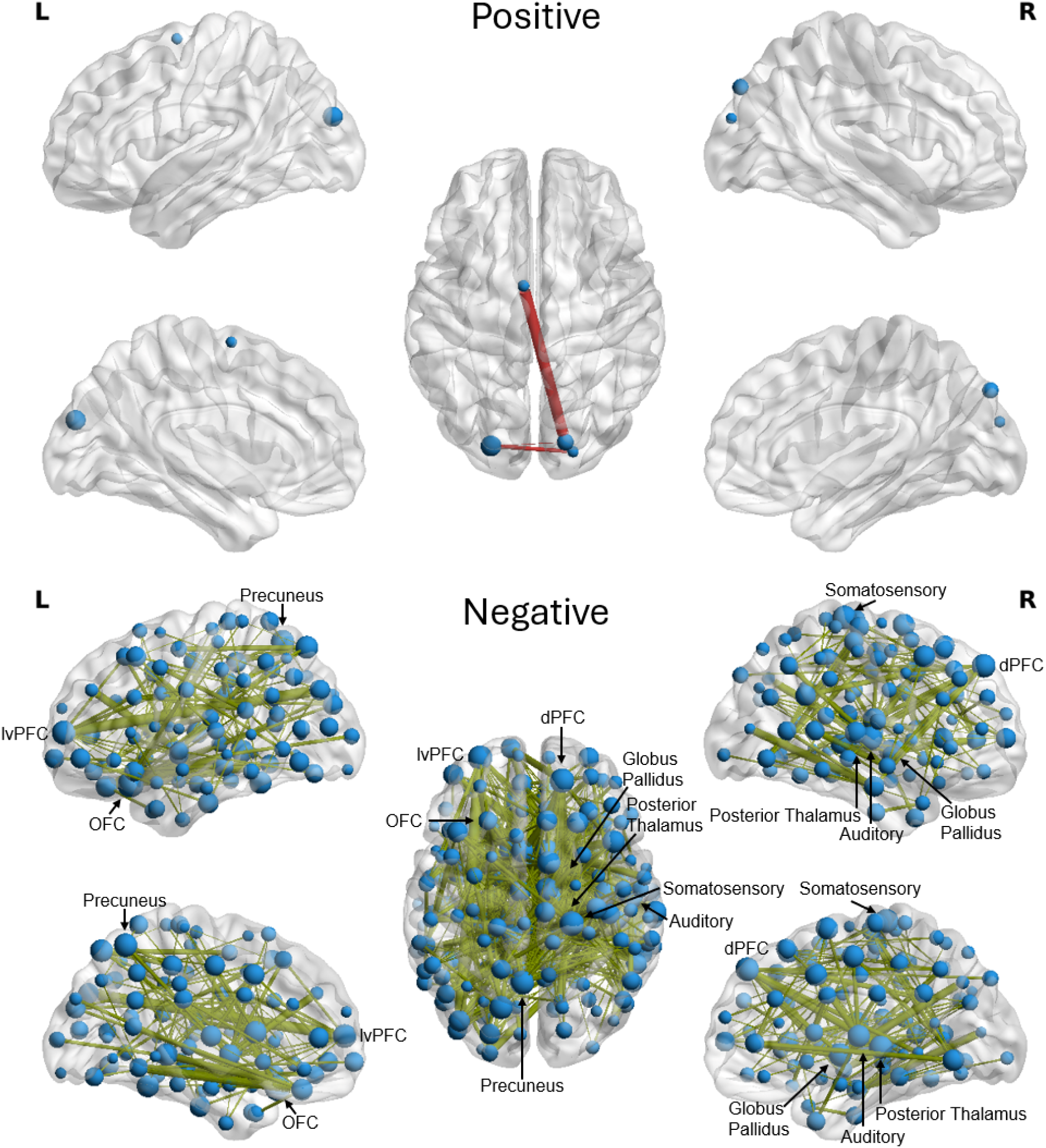
Functional connectivity cluster identified by NBS analysis, representing a significant difference between YA and OA on the sleep difference network (restricted minus normal sleep). Connection size indicates the magnitude of the t-score. Highest degree regions are labelled (lvPFC: lateral ventral prefrontal cortex; OFC: orbitofrontal cortex; dPFC: dorsal prefrontal cortex).

An additional mean-centred PLS was conducted on the functional connectivity with the effect of sex additionally included. One significant LV was identified (*p* = .015), with functional connections showing a similar pattern of association with sleep and age as in the previous PLS analysis, while also identifying a sex effect, in which a sleep effect is observed in these connections for males but not for females (see Figure 5). This is consistent with the pattern of the individual brain scores that we observed for the previous PLS, showing that the females were not consistently following the pattern identified by the LV. The functional connections identified as reliably associated with the LV formed a network with highly connected regions, including the LH caudate and RH auditory cortex, as observed in the previous PLS and NBS analyses, as well as the LH temporal occipital cortex, inferior parietal lobule (IPL), RH secondary somatosensory cortex, and inferior parietal sulcus (IPS; see Figure 6). Of the 1109 identified connections, 27.054% were LH connections, 24.464% were RH connections, and 48.482% were interhemispheric connections.

**Figure 5.**
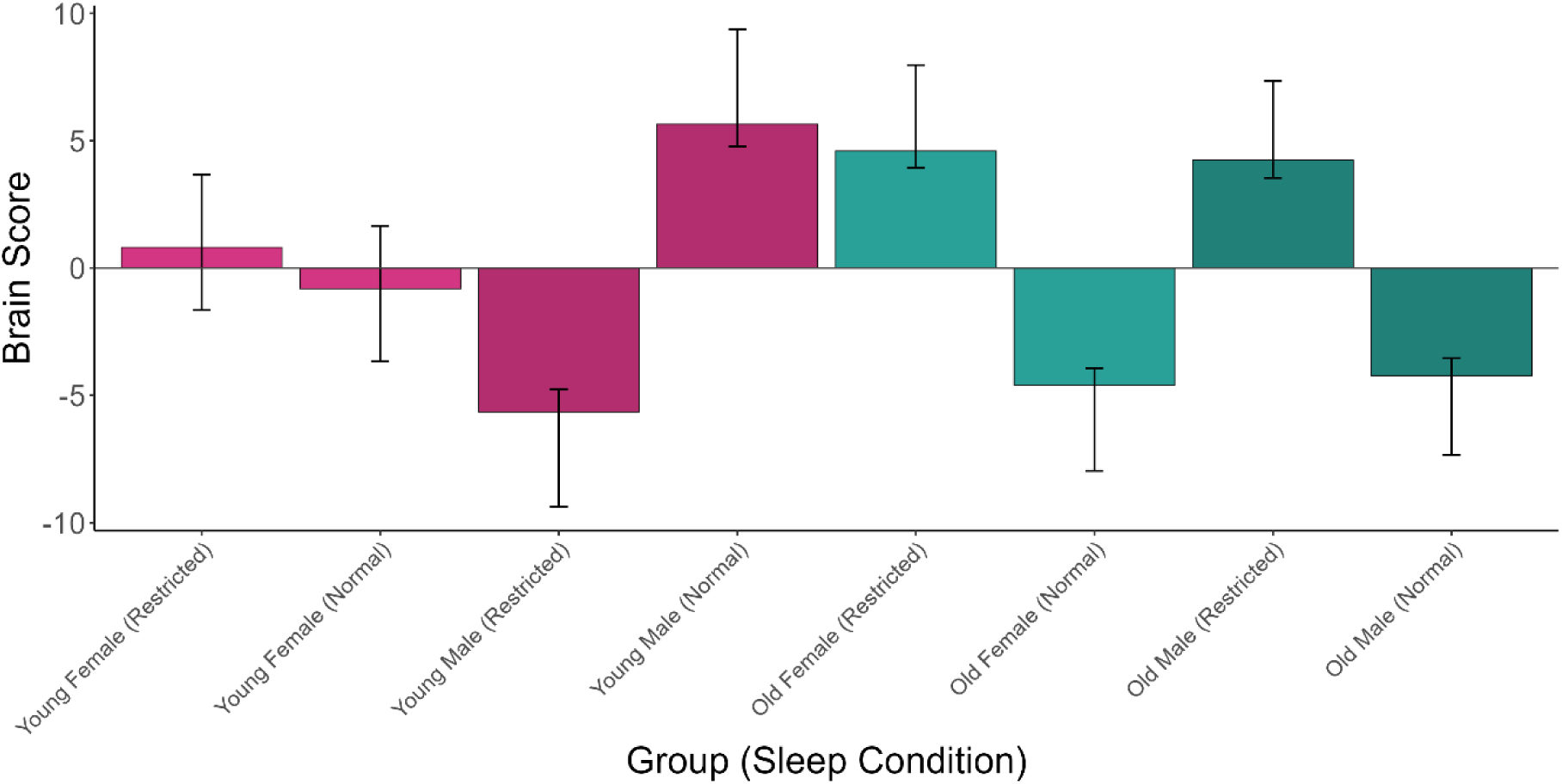
Mean brain scores from FC mean-centred PLS LV, for participants in the young and old age groups separated by sex, in restricted and normal sleep conditions. Error bars represent 95% confidence intervals.

**Figure 6.**
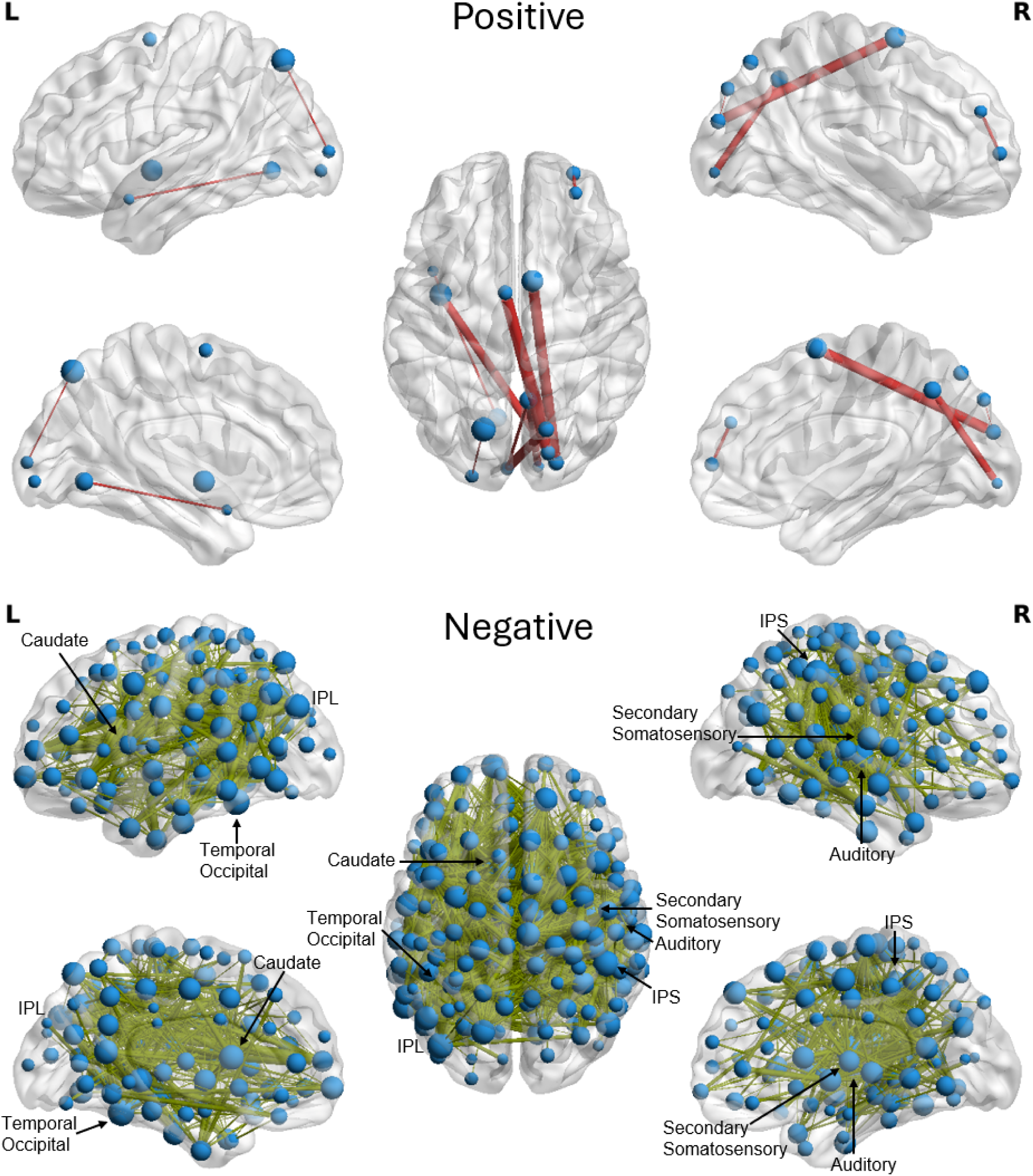
Reliable bootstrap ratios from FC mean-centred PLS LV with independent variables of age, sex, and sleep condition, depicted separately for positive and negative values. Connection size indicates the magnitude of the BSR. Highest degree regions are labelled (IPL: inferior parietal lobule; IPS: inferior parietal sulcus).

### Functional Connectivity Degree

A mean-centred PLS analysis was performed with FC degree (total regional connectivity weights) as the dependent variable and independent variables of age group and sleep condition. This analysis identified one significant LV (*p* = .021) associated with a crossover interaction between age and sleep, whereby the effect of sleep restriction occurred in opposite directions for YA and OA (see Figure 7A). As seen by the lines showing the sleep effect for individuals by sex, for the YA males, all but three of these individuals followed the pattern identified by the LV, while YA females were much more inconsistent. For OA, all but two of the males followed the pattern identified by the LV, and all but two of the females followed the pattern (see Figure 7A). The functional connections reliably associated with this LV were all negatively associated, indicating that sleep restriction in OA was associated with reduced degree in these regions, but increased degree in YA (see Figure 7B). The pattern of cortical regions identified by this LV resembles the association end of the Sensorimotor-Association (S-A) Axis (e.g., Luo et al., 2024; see Figure 7C), and indeed, the cortical BSR values from this LV were negatively correlated with the S-A Axis values, *R*(218) = -0.362, *p* < .001. Looking at the mean degree difference between restricted and normal sleep for these regions, OA exhibited a negative difference for all regions and YA exhibited a positive difference for these regions, consistent with the pattern identified by the LV (see Figure 8).

**Figure 7.**
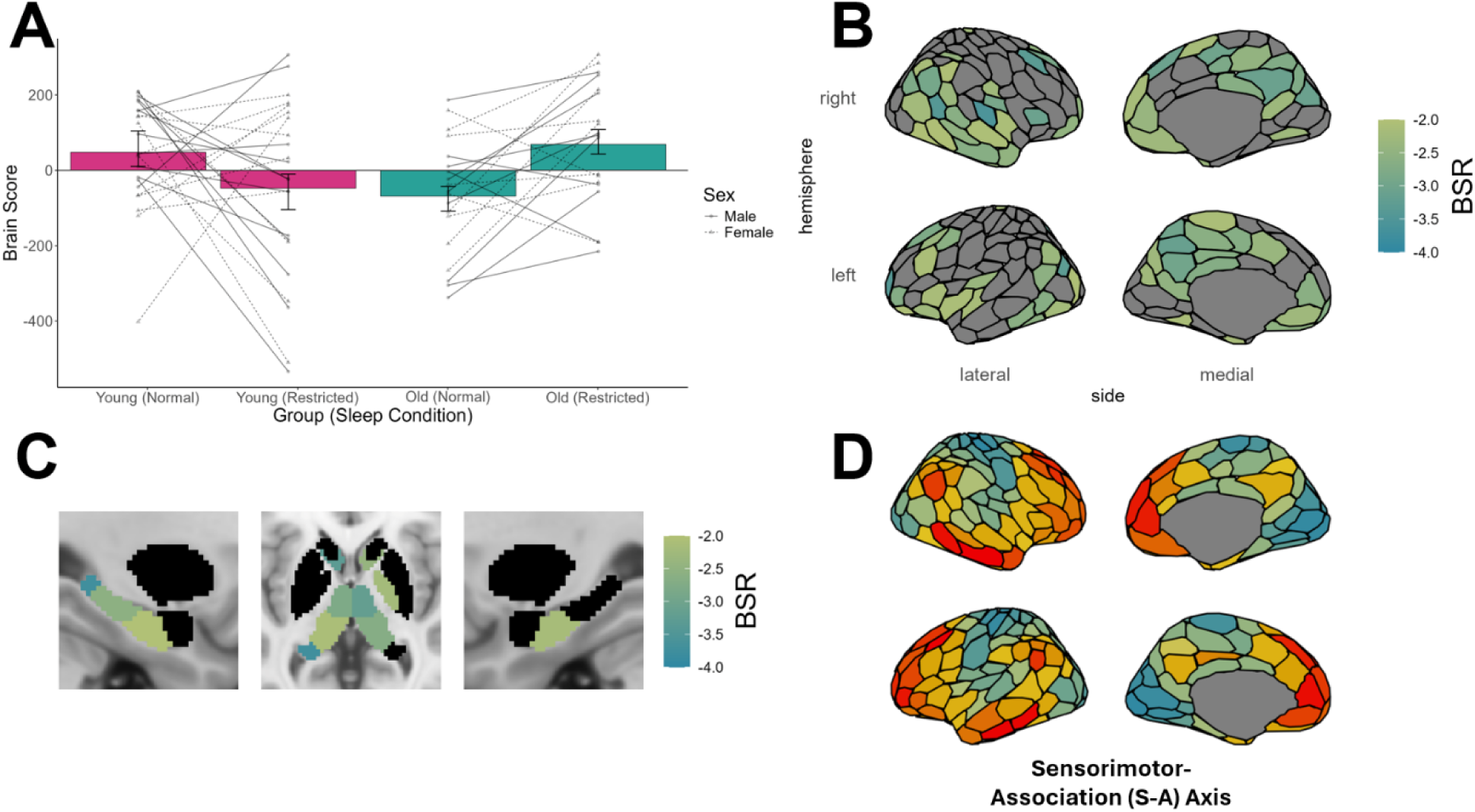
**A)** Mean brain scores from FC degree mean-centred PLS LV, for participants in the young and old age groups, in restricted and normal sleep conditions. Error bars represent 95% confidence intervals. Solid (male) and dashed (female) lines depict the change in the individual brain score values between sleep conditions. **B)** Reliable cortical bootstrap ratios from FC degree mean-centred PLS LV with independent variables of age and sleep condition. **C)** Reliable subcortical bootstrap ratios from FC degree mean-centred PLS LV with independent variables of age and sleep condition. **D)** For comparison, the Sensorimotor-Association Axis (from blue/green sensorimotor regions to yellow/red association regions) from Luo et al. (2024). Regions identified as reliable overlap with the association end of the S-A axis and BSRs are negatively correlated with the S-A axis values, *R*(218) = -0.362, *p* < .001.

**Figure 8.**
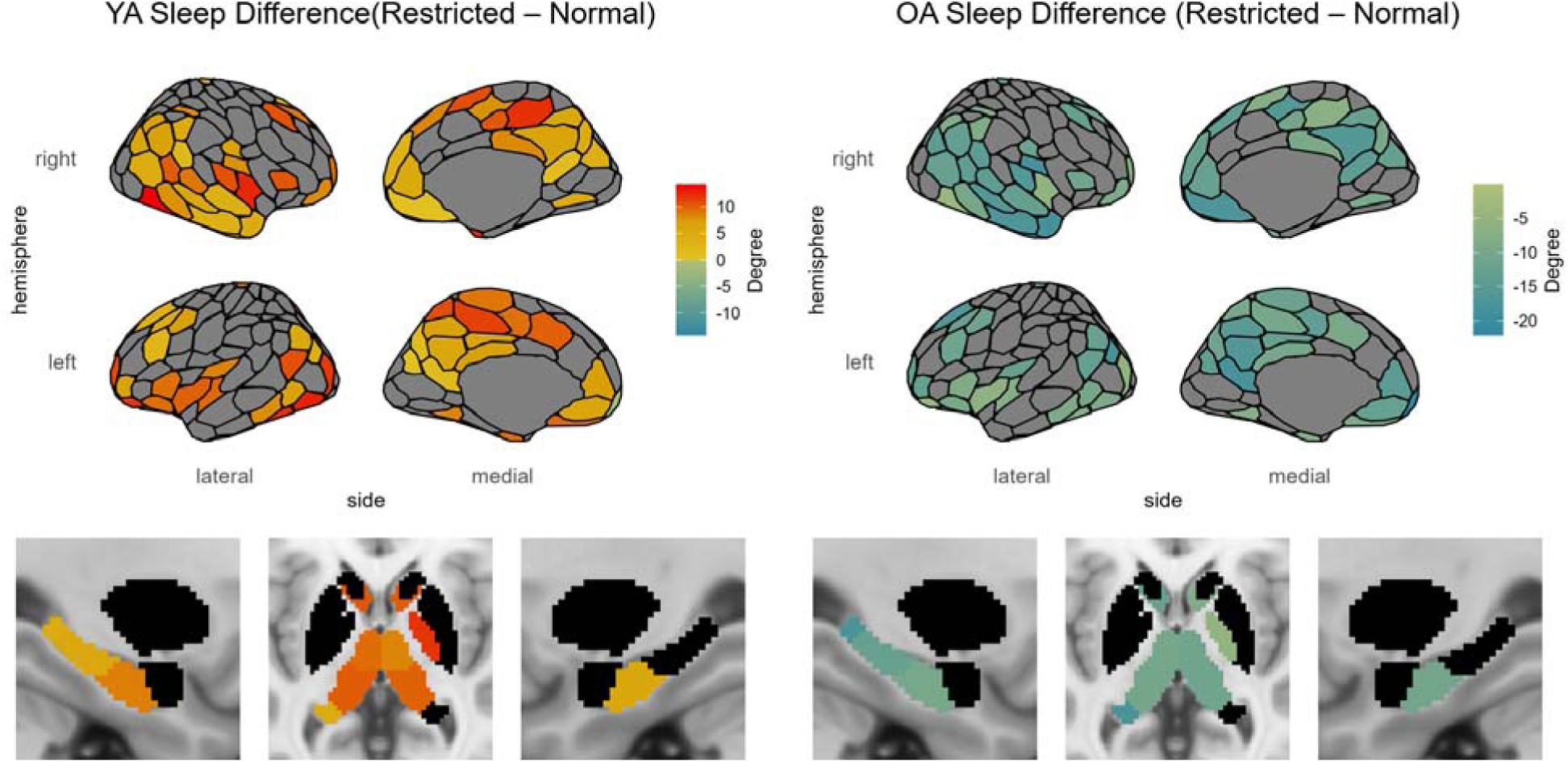
Mean FC degree difference between restricted and normal sleep conditions in YA (left) and OA (right) for regions identified as reliable by the FC degree mean-centred PLS LV.

An additional mean-centred PLS was conducted on the functional connectivity degree with the effect of sex additionally included. One significant LV was identified (*p* = .004), with degree showing a similar pattern of association with sleep and age as in the previous PLS analysis, while also identifying a sex effect, in which a sleep effect is observed in these connections for males but not for females (see Figure 9). This is consistent with the pattern of the individual brain scores that we observed for the previous PLS, showing that the females were not consistently following the pattern identified by the LV. The regions identified as reliably associated with the LV were consistent with those observed in the previous PLS analysis, but with additional regions identified (see Figure 10).

**Figure 9.**
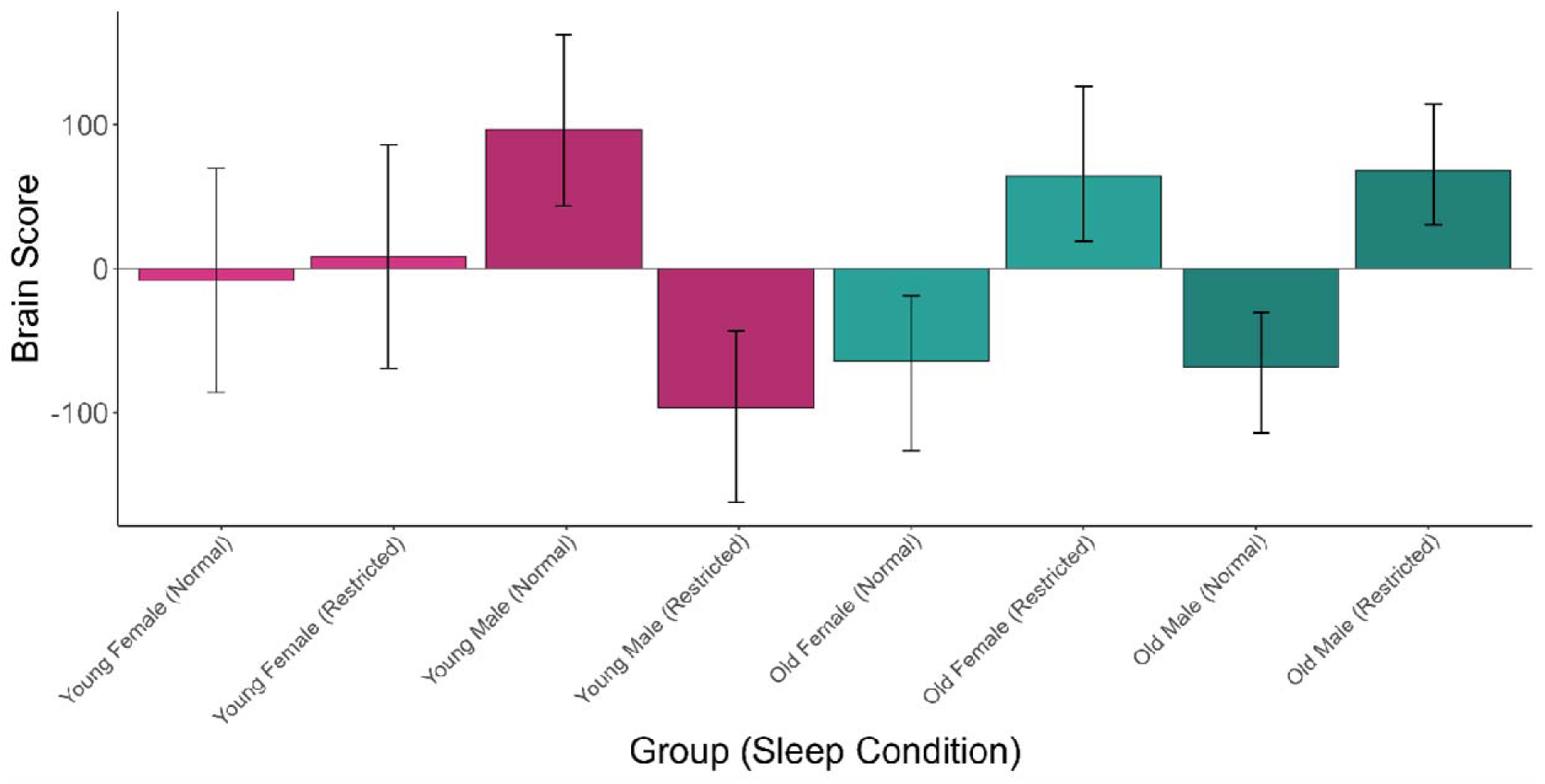
Mean brain scores from FC degree mean-centred PLS LV, for participants in the young and old age groups separated by sex, in restricted and normal sleep conditions. Error bars represent 95% confidence intervals.

**Figure 10.**
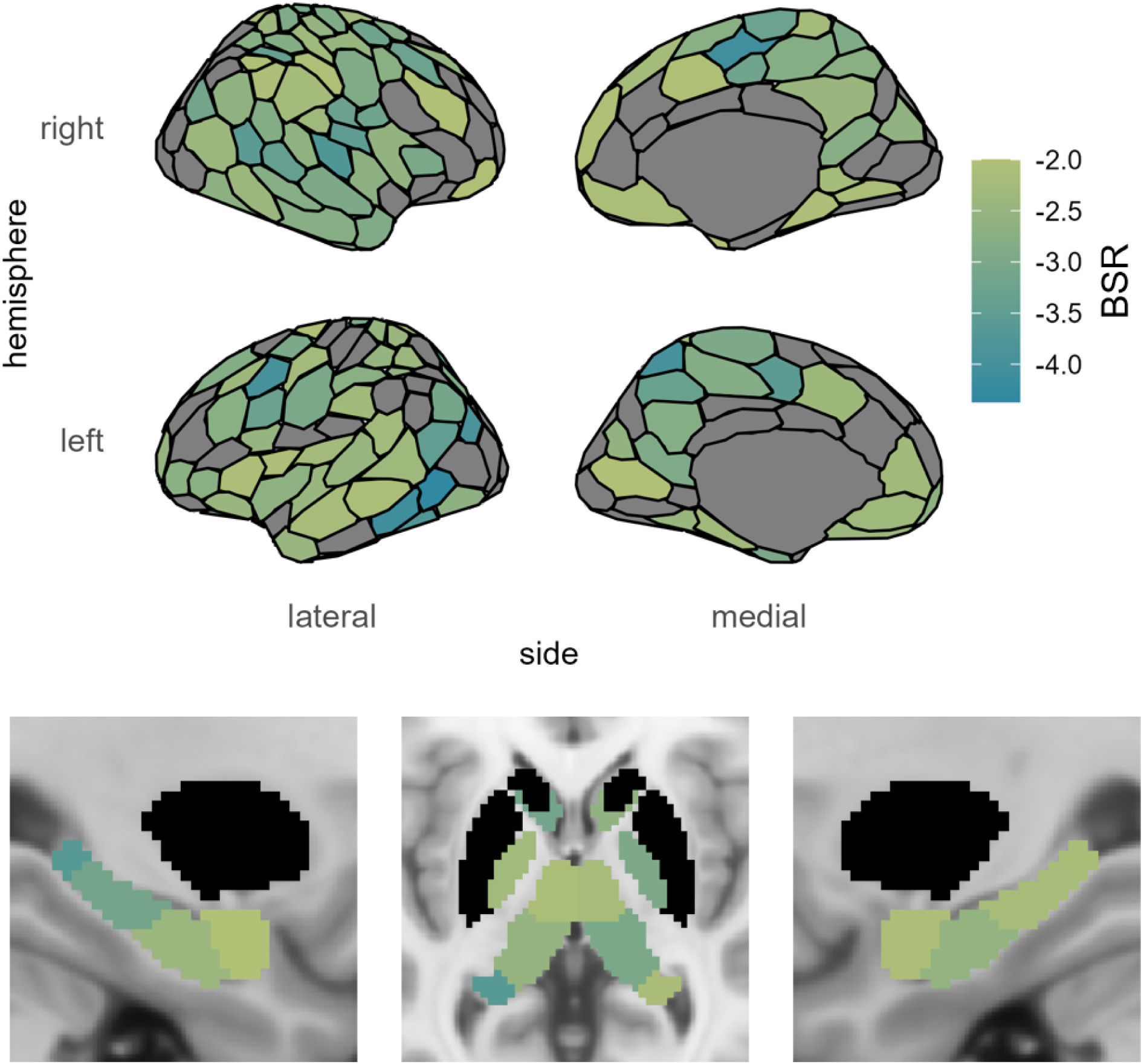
Reliable bootstrap ratios from FC degree mean-centred PLS LV with independent variables of age, sex, and sleep condition.

### Dynamic Functional Connectivity

The LEiDA dFC analysis identified 5 states, interpreted as Global Coherence, Default Mode Network (DMN), Frontoparietal Network, Somatormotor, and Attention states. The fractional occupancy (FO) of these states (relative time spent in each state) was analyzed using a mean-centred PLS with FO as the dependent variable and independent variables of age group and sleep condition. One significant LV (*p* = .018) was identified, exhibiting the same cross-over interaction association as shown previously (see Figure 11A). This LV identified a decrease in the amount of time spent in the Global Coherence state for OA after restricted sleep, and an opposite effect of sleep restriction on YA (see Figure 11B for BSRs and Figure 11C for mean FO of the Global Coherence state). In the YA group, 5/12 males and 6/11 females followed the pattern of association identified by the LV, and in the OA group, 8/10 males and 8/9 females followed the pattern of association identified by the LV (see Figure 11A).

**Figure 11.**
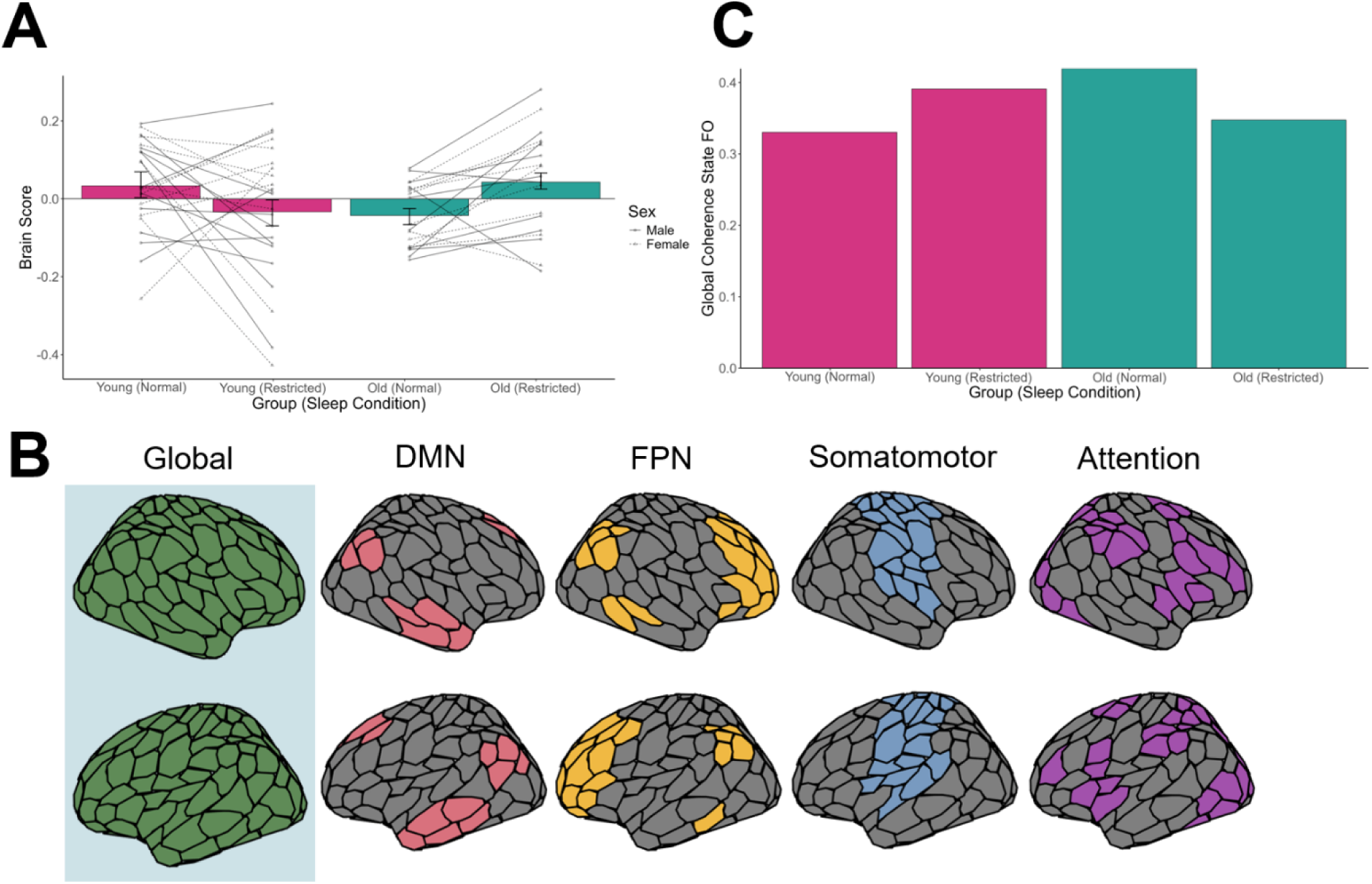
**A)** Mean brain scores from LEiDA FO mean-centred PLS LV, for participants in the young and old age groups, in restricted and normal sleep conditions. Error bars represent 95% confidence intervals. Solid (male) and dashed (female) lines depict the change in the individual brain score values between sleep conditions. **B)** Reliable bootstrap ratios from LEiDA FO mean-centred PLS LV with independent variables of age and sleep condition (blue highlighting indicates a significant negative BSR for the Global Coherence State). **C)** Mean FO of the Global Coherence State in each of the conditions.

An additional PLS analysis including sex as an independent variable did not produce a significant LV. An analysis of the transition probability matrix, representing the likelihood of each possible state transition, was analyzed using a mean-centred PLS with independent variables of age and sleep, but produced no significant LV.

### Network Modularity

A network modularity analysis was conducted based on functional connectivity modules identified by the Louvain algorithm. High modularity indicates that the network has strong connectivity within modules but weaker connectivity between modules. A linear mixed effects (LMM) model identified a significant crossover interaction between age and sleep, whereby modularity increased in the restricted sleep condition for OA but decreased for OA (see Figure 12 and Table 1). Although age was not significant in the LMM, the 95% confidence intervals indicate that OA had less modularity than YA in the normal sleep condition, replicating past research (Song et al., 2014), and that under restricted sleep conditions YA modularity was reduced to normal OA levels, replicating others (Zhou et al., 2017). A decrease in YA modularity has also been demonstrated in other research (Ben Simon et al., 2017). Additionally, we demonstrated that OA modularity is increased to normal YA levels in the restricted sleep condition, representing a novel discovery. An additional model was investigated with sex added as an independent variable, but sex was not significant and did not alter the findings of the previous model. A supplementary LMM was conducted with the attention score from the BADD ADHD assessment added as an independent variable. In this model, the age × sleep interaction was still significant (*t* = 2.890), but the attention measure did not reach significance (*t* = 1.813). However, the attention measure is trending in a positive direction, and if significant would suggest that the increased modularity after restricted sleep may represent a positive network alteration for OA. Other research has positively associated modularity with cognitive performance (Bertolero et al., 2018), so this possibility should be investigated in future research.

**Figure 12.**
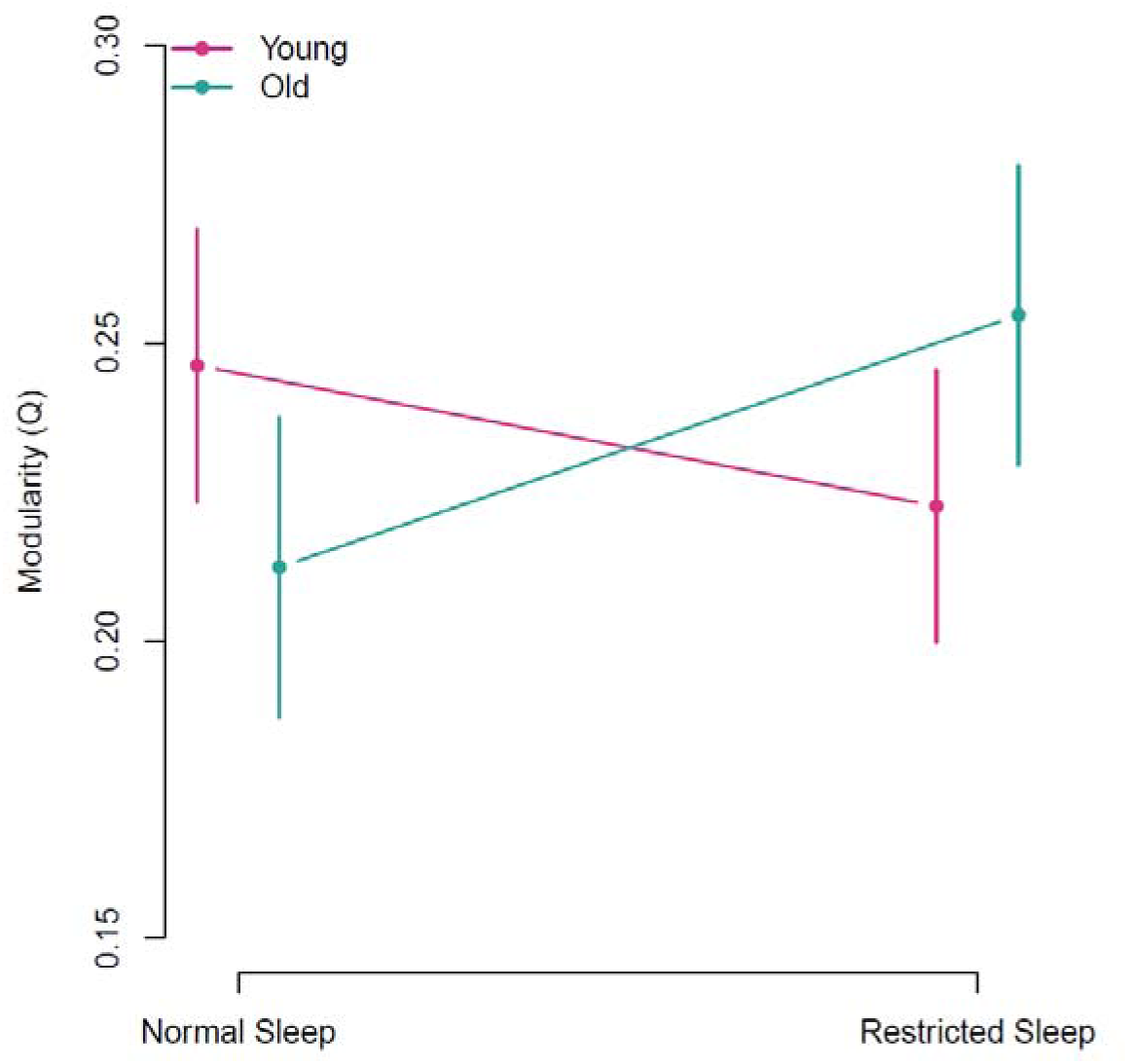
FC modularity as a function of age and sleep. Error bars represent the 95% confidence interval.

**Table 1.** Linear Mixed Model results with modularity as the dependent variable and independent variables age and sleep plus their interaction, including a random intercept for participants.

|  | Errors |  |  |
| --- | --- | --- | --- |
| Random Effects | Variance | SD |  |
| **Participants** |  |  |  |
| Intercept | 6.291×10<sup>-4</sup> | .025 |  |
| Fixed Effects | Estimate (OR) | Std. Error | t-value |
| Intercept | .246 | .012 | 21.417 |
| Sleep (Restricted) | -.024 | .014 | -1.633 |
| Age (Old) | -.034 | .017 | -1.985 |
| Sleep (Restricted) × Age (Old) | .066 | .022 | *3.067 |

### Signal Variability

In a departure from the rest of the analyses presented here, we analyzed a measure not based on network data: grey matter signal variability. This analysis serves primarily as a replication attempt of the main significant finding from the authors that produced this dataset (Nilsonne et al., 2017). We included independent variables of age, sleep, and motion as well as the age × sleep in a linear mixed effects model with a random intercept of participants and dependent variable log-transformed grey matter signal variability (motion was included for consistency with the previous research). We were able to replicate the age effect demonstrated previously, but not the age × sleep interaction (see Figure 13 and Table 2). Especially when it comes to global signal measures like signal variability, the preprocessing steps can make a difference in the final results (Liu et al., 2017). Given that the original research on this dataset utilized very different preprocessing (largely SPM instead of FSL), we expect this may account for the difference in findings. An additional model was investigated with sex added as an independent variable, but sex was not significant and did not alter the findings of the previous model.

**Figure 13.**
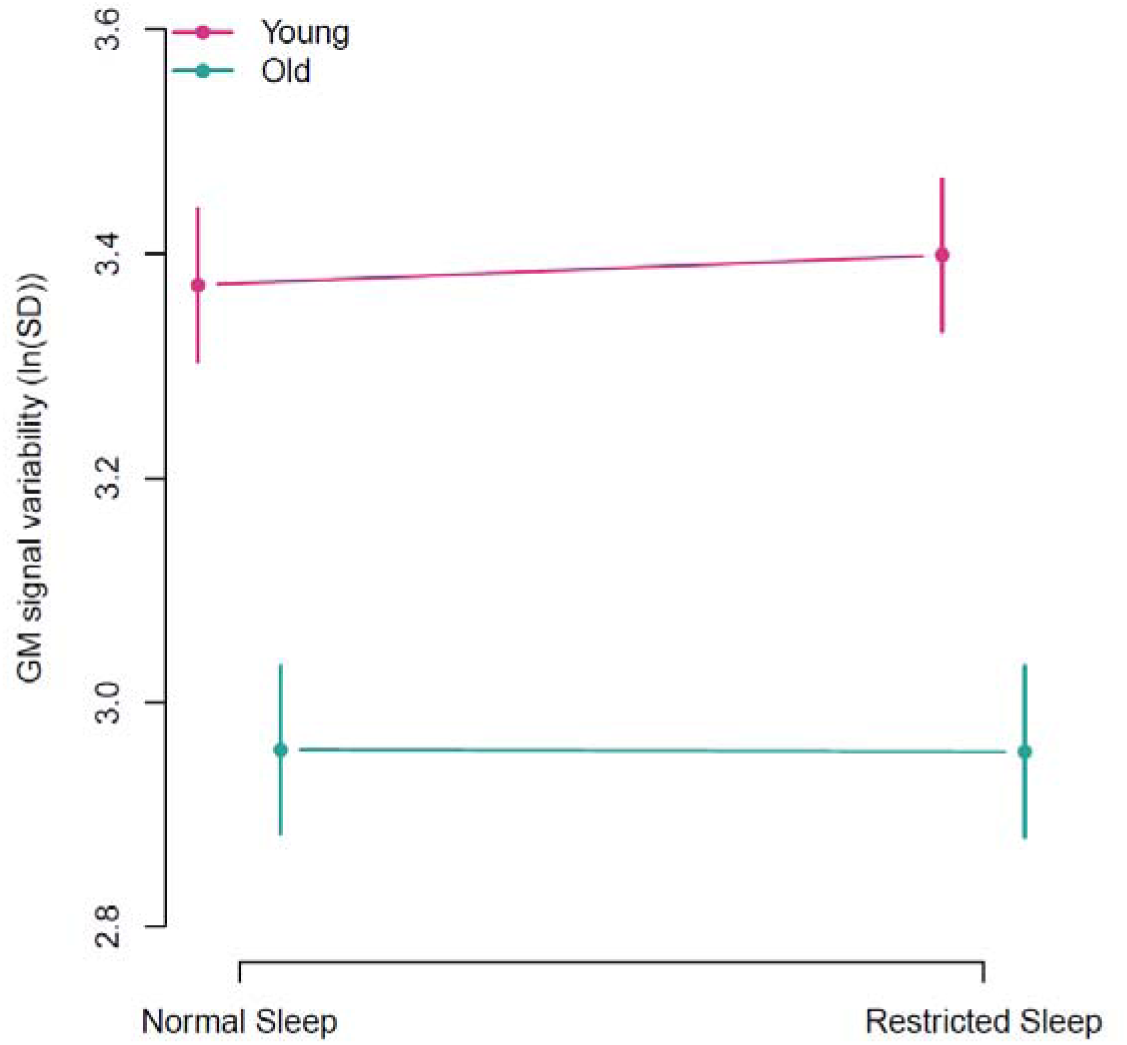
Grey matter signal variability (log-transformed standard deviation of signal) as a function of age and sleep. Error bars represent the 95% confidence interval.

**Table 2.** Linear Mixed Model results with log-transformed GM signal variability as the dependent variable and independent variables age, sleep, and motion, plus the age × sleep interaction, including a random intercept for participants.

|  | Errors |  |  |
| --- | --- | --- | --- |
| Random Effects | Variance | SD |  |
| **Participants** |  |  |  |
| Intercept | 4.226×10<sup>-3</sup> | .065 |  |
| Fixed Effects | Estimate (OR) | Std. Error | t-value |
| Intercept | 3.386 | .041 | 81.706 |
| Sleep (Restricted) | .027 | .044 | .608 |
| Age (Old) | -.414 | .051 | *-8.198 |
| Sleep (Restricted) × Age (Old) | -.028 | .066 | -.428 |
| Motion | -.040 | .073 | -.551 |

### Voxelwise Functional Connectivity

Multivariate distance matrix regression (MDMR) analyses were performed on the voxelwise functional connectivity matrices comparing sleep conditions within age groups, and on the voxelwise functional connectivity sleep difference matrices (subtracting the connectivity values for normal sleep from those obtained in the restricted sleep condition) between age groups. Of these analyses, only the sleep condition comparison within YA identified significant voxels, identifying a cluster of voxels primarily localized within the bilateral medial, orbitofrontal, and dorsolateral PFC (see Figure 14). Although we were unable to detect a voxelwise sleep-related difference in FC between age groups with this dataset, the granular level of detail about the network effect of acute sleep restriction in YA has not been reported in the literature and represents an important contribution to network neuroscience sleep research.

**Figure 14.**
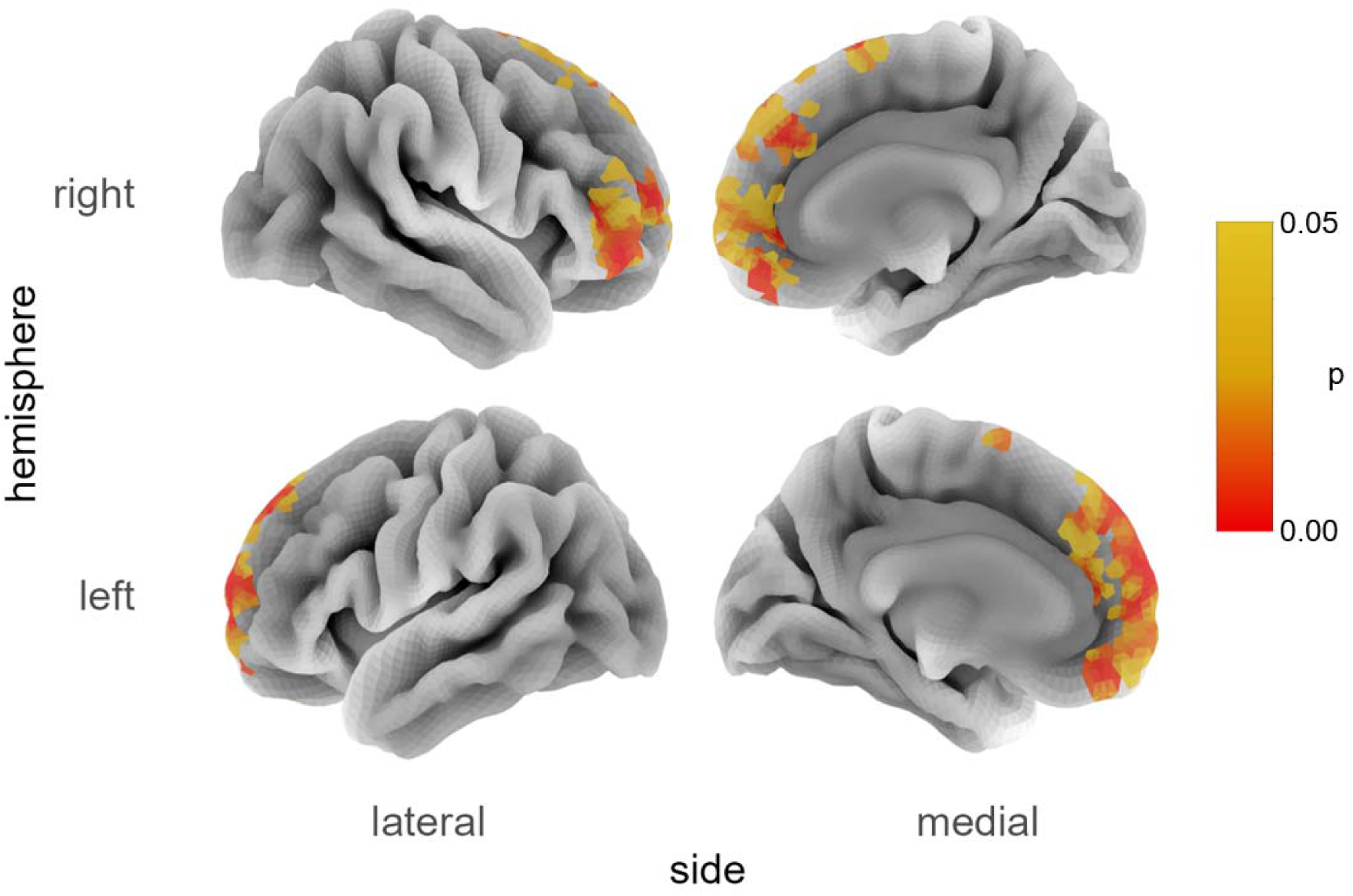
Voxels exhibiting a sleep condition dependent difference in their FC connectivity profile as identified by multivariate distance matrix regression for YA.

## Discussion

We have identified multiple aspects of the functional connectivity network that exhibit opposite effects of acute sleep restriction when comparing older and younger adults. Both mean-centred PLS and NBS methods identified a significant subnetwork associated with sleep restriction, composed primarily of connections that were weaker after sleep restriction for older adults but, conversely, were stronger after sleep restriction for younger adults. The most highly implicated regions within this subnetwork included LH OFC and RH dPFC, somatosensory cortex, auditory cortex, and posterior thalamus, which both PLS and NBS identified. Using graph theory to investigate the regional network effects of sleep restriction, we found a large number of regions where functional connectivity degree decreased after sleep restriction for older adults but increased after sleep restriction for younger adults. The regions identified were primarily composed of association regions. Sex-based analyses of both the edge- and region-focused analyses revealed that males drove the effects seen in younger adults. In comparison, the effect in older adults was common across both sexes. A dynamic functional connectivity analysis identified 5 states, their fractional occupancies, and their transition probabilities. Older adults spent less time (measured via fractional occupancy) in the global coherence state after sleep restriction, while younger adults spent more time in this state after sleep restriction. Furthermore, functional connectivity modularity also demonstrated a crossover interaction with age and sleep, whereby younger adults exhibited reduced modularity after sleep restriction. For younger adults, modularity fell to the level of older adults under normal sleep conditions, while older adults exhibited enhanced modularity after sleep restriction, boosted to levels corresponding to younger adults under normal sleep conditions. While we were not able to identify age differences in voxelwise functional connectivity profiles using MDMR, we did identify a cluster of significant voxels exhibiting a sleep difference for younger adults, localized to the bilateral medial PFC, orbitofrontal cortex, and dorsolateral PFC.

These findings represent a novel demonstration of a consistent cross-over interaction between the effects of age and acute sleep loss on brain network measures. Finding an interaction of these variables has been attempted previously without success (Nilsonne et al., 2017), but we were able to observe the interaction across numerous metrics by focusing on edge- and region-centric analyses, which allow for the network to be inspected in finer detail. Our dynamic functional connectivity analyses also represent a novel investigation in this domain, which produced a consistent interaction when looking at state fractional occupancy. We also discovered that the whole-brain metric of modularity exhibited the same cross-over interaction, extending past research that has looked at healthy young adults in normal vs. deprived sleep conditions as well as older adults in a normal sleep condition, but lacked the fourth condition of sleep deprived older adults needed to identify this interaction (Zhou et al., 2017). With all four of these conditions, we were able to replicate past findings that older adults have reduced modularity under normal sleep conditions and that younger adults exhibit reduced modularity in the range of older adult default levels, while demonstrating the novel finding that older adults under sleep restricted conditions have modularity levels that are akin to well-rested young adults. Given that modularity is associated with better cognitive performance (Bertolero et al., 2018), these findings suggest that acute sleep restriction may induce compensatory reorganization of functional connectivity for older adults. Compensatory reactions of the brain to sleep restriction would not be surprising, given the past research on young adults that has demonstrated brain function compensation in response to sleep loss (Caldwell et al., 2004; Chengyang et al., 2017; Drummond et al., 2000; Tian et al., 2024).

The idea that *acute* loss of sleep may result in temporary positive effects may seem counterintuitive, given what we know about the negative effects of *chronic* loss of sleep. However, the chronic effects of sleep loss in older age are a cumulative effect across the lifespan, which has the greatest impact in younger adulthood and middle age, increasing the risk of dementia and cognitive decline, whereas sleep loss in older adults does not impact risk of cognitive decline or dementia consistently or to the same extent as earlier in the lifespan (as reviewed by Scullin & Bliwise, 2015). Furthermore, there are positive features of acute sleep loss that have been observed in previous research. For example, although chronic lack of sleep is associated with increased depression symptoms (Fang et al., 2019; Krakow et al., 2000), acute sleep deprivation can temporarily reduce symptoms of depression in around half of depression and posttraumatic stress disorder patients (Boland et al., 2017; Germain et al., 2008; Hemmeter et al., 1998, 2010).

It may also be surprising, at first, that older adult brain networks consistently respond to sleep restriction in an opposite way to young adults. However, research has demonstrated that cognitive abilities are negatively affected by sleep restriction for younger adults, while older adults are relatively or completely spared from these effects (Duffy et al., 2009; Gerhardsson et al., 2019; Scullin & Bliwise, 2015). Similarly, the benefits of napping on cognitive function are often only seen for young and middle-aged adults, and the benefits of sleep for memory consolidation are stronger for younger adults than older adults (Scullin & Bliwise, 2015).

Additionally, recent research from our lab has shown multiple ways in which the healthy aging brain network reorganizes to support cognitive ability, utilizing a distinct network regime that is different in fundamental ways from the healthy young adult brain (Heisz et al., 2015; Neudorf et al., 2024, 2025). For these reasons, it may be expected that the way the young and old adult brain networks adapt to restricted sleep are also fundamentally different in the ways we have identified here.

Our results provide insight into theories that have been developed to explain the discrepancy between younger and older adults in the effects of sleep restriction on cognitive abilities. As we have highlighted, older adults are relatively or completely spared from the effects of sleep restriction, unlike younger adults (Scullin & Bliwise, 2015). Our findings support the consolidation theory most strongly, as we did not simply observe in older adults the absence or decrease of a sleep restriction effect seen for younger adults, as would be expected based on the reduced sleep need and functional weakening theories. Rather, we consistently identified brain network features that changed after sleep restriction in opposite directions for younger and older adults. This finding is consistent with the theory that the brain compensates for changes due to age by reorganizing the brain in a fundamental way.

## Limitations and Future Directions

The dataset used for this analysis did not include extensive cognitive assessment measures, and did not include measurements in both sleep conditions to assess the impact of sleep restriction on cognitive performance. For this reason, the analyses we conducted here should be replicated using data that includes cognitive performance measures pre- and post- restricted sleep, in order to determine which of the brain network responses to acute sleep restriction may be compensatory.

Furthermore, a larger sample size with more variation in age (e.g., filling in the gap between 30 – 65 years old that is missing from the current research) would help to increase the power of these analyses and allow for detection of more subtle age effects. In particular, looking at middle age and the transition to older adulthood could uncover how the brain’s response to sleep restriction transitions over time into such opposite patterns of sleep restriction-induced reorganization.

Additionally, including an analysis of biomarkers for dementia (e.g., amyloid, tau, alpha-synuclein, and neurodegeneration) would be beneficial, particularly because sleep disturbances are risk factors and symptoms of several distinct neurodegenerative diseases. For this reason, these biomarkers could confound the pure effects of age and should be included in future research.

## Conclusions

We have demonstrated yet another way in which the old adult brain represents a distinct brain network regime from the young adult brain. Younger and older adults’ brain networks respond to low sleep conditions in opposite ways, as measured using functional connectivity, degree, modularity, and dynamic functional connectivity state fractional occupancy. The opposing effects were observed based on these multiple brain network variables, suggesting a consistent and fundamental difference in the effects of sleep restriction on younger and older adults. Further research should continue to investigate these effects and how they relate to preserved cognitive ability and brain health.

## Data and Code Availability

Data were downloaded from OpenfMRI (https://openfmri.org/dataset/ds000201/).

Code used to produce analyses available via github repository (https://github.com/McIntosh-Lab/SleepyBrain_analyses).

## Ethics Statement

Details about recruitment, ethics, and the full dataset can be obtained from the original dataset paper and a follow-up analysis paper (Nilsonne et al., 2016, 2017).

## Author Contributions

Josh Neudorf contributed to Conceptualization, Data curation, Formal analysis, Investigation, Methodology, Software, Validation, Visualization, Writing – original draft, and Writing – review & editing.

Leanne Rokos contributed to Data curation, Methodology, Validation, and Writing – review & editing.

Kelly Shen contributed to Conceptualization, Data curation, Investigation, Methodology, Project administration, Resources, Validation, and Writing – review & editing.

Brianne Kent contributed to Methodology, Validation, and Writing – review & editing.

Anthony R. McIntosh contributed to Conceptualization, Data curation, Funding acquisition, Investigation, Methodology, Project administration, Resources, Supervision, Validation, and Writing – review & editing.

## Acknowledgments

This research was supported by the Natural Sciences and Engineering Research Council of Canada (NSERC) Postdoctoral Fellowships Program and by the Hilary and Galen Weston Foundation through Canadian Neuroparalytic Scholars Program funding to Josh Neudorf, and by NSERC Discovery Grant RGPIN-2018-04457 and Canadian Institutes of Health Research (CIHR) Project Grant PJT-168980 to the senior author Anthony R. McIntosh. We thank Jaspreet Dodd for her assistance in validating the fMRI data preprocessing. This research was enabled in part by support provided by the British Columbia DRI Group and the Digital Research Alliance of Canada (alliancecan.ca). The authors affirm that there are no conflicts of interest to disclose.

## References

Bateman, R. J., Xiong, C., Benzinger, T. L. S., Fagan, A. M., Goate, A., Fox, N. C., Marcus, D. S., Cairns, N. J., Xie, X., Blazey, T. M., Holtzman, D. M., Santacruz, A., Buckles, V., Oliver, A., Moulder, K., Aisen, P. S., Ghetti, B., Klunk, W. E., McDade, E., … Dominantly Inherited Alzheimer Network. (2012). Clinical and biomarker changes in dominantly inherited Alzheimer’s disease. The New England Journal of Medicine, 367(9), 795–804. 10.1056/NEJMoa1202753

Ben Simon, E., Maron-Katz, A., Lahav, N., Shamir, R., & Hendler, T. (2017). Tired and misconnected: A breakdown of brain modularity following sleep deprivation. Human Brain Mapping, 38(6), 3300–3314. 10.1002/hbm.23596

Boland, E. M., Rao, H., Dinges, D. F., Smith, R. V., Goel, N., Detre, J. A., Basner, M., Sheline, Y. I., Thase, M. E., & Gehrman, P. R. (2017). Meta-Analysis of the Antidepressant Effects of Acute Sleep Deprivation. The Journal of Clinical Psychiatry, 78(8), e1020– e1034. 10.4088/JCP.16r11332

Buckner, R. L., Krienen, F. M., Castellanos, A., Diaz, J. C., & Yeo, B. T. T. (2011). The organization of the human cerebellum estimated by intrinsic functional connectivity. Journal of Neurophysiology, 106(5), 2322–2345. 10.1152/jn.00339.2011

Cabral, J., Vidaurre, D., Marques, P., Magalhães, R., Silva Moreira, P., Miguel Soares, J., Deco, G., Sousa, N., & Kringelbach, M. L. (2017). Cognitive performance in healthy older adults relates to spontaneous switching between states of functional connectivity during rest. Scientific Reports, 7(1), Article 1. 10.1038/s41598-017-05425-7

Caldwell, J. A., Smith, J. K., Caldwell, J. L., Mu, Q., George, M., Peters, G., & Brown, D. L. (2004). *Functional Magnetic Resonance Imaging Shows Potential for Predicting Individual Differences in Fatigue Vulnerability: (425102005-001)* [Dataset]. 10.1037/e425102005-001

Caliński, T., & Harabasz, J. (1974). A dendrite method for cluster analysis. Communications in Statistics, 3(1), 1–27. 10.1080/03610927408827101

Chee, M. W. L., & Zhou, J. (2019). Chapter 7—Functional connectivity and the sleep-deprived brain. In H. P. A. Van Dongen, P. Whitney, J. M. Hinson, K. A. Honn, & M. W. L. Chee (Eds.), Progress in Brain Research (Vol. 246, pp. 159–176). Elsevier. 10.1016/bs.pbr.2019.02.009

Chengyang, L., Daqing, H., Jianlin, Q., Haisheng, C., Qingqing, M., Jin, W., Jiajia, L., Enmao, Y., Yongcong, S., & Xi, Z. (2017). Short-term memory deficits correlate with hippocampal-thalamic functional connectivity alterations following acute sleep restriction. Brain Imaging and Behavior, 11(4), 954–963. 10.1007/s11682-016-9570-1

Cirelli, C. (2012). Brain Plasticity, Sleep and Aging. Gerontology, 58(5), 441–445. 10.1159/000336149

Cohen, J. R. (2018). The behavioral and cognitive relevance of time-varying, dynamic changes in functional connectivity. NeuroImage, 180, 515–525. 10.1016/j.neuroimage.2017.09.036

Dai, C., Zhang, Y., Cai, X., Peng, Z., Zhang, L., Shao, Y., & Wang, C. (2020). Effects of Sleep Deprivation on Working Memory: Change in Functional Connectivity Between the Dorsal Attention, Default Mode, and Fronto-Parietal Networks. Frontiers in Human Neuroscience, 14, 360. 10.3389/fnhum.2020.00360

Dautricourt, S., Gonneaud, J., Landeau, B., Calhoun, V. D., de Flores, R., Poisnel, G., Bougacha, S., Ourry, V., Touron, E., Kuhn, E., Demintz-King, H., Marchant, N. L., Vivien, D., de la Sayette, V., Lutz, A., Chételat, G., Arenaza-Urquijo, E. M., Allais, F., André, C., … for the Medit-Ageing Research Group. (2022). Dynamic functional connectivity patterns associated with dementia risk. Alzheimer’s Research & Therapy, 14(1), 72. 10.1186/s13195-022-01006-7

Deco, G., Cabral, J., Woolrich, M. W., Stevner, A. B. A., van Hartevelt, T. J., & Kringelbach, M. L. (2017). Single or multiple frequency generators in on-going brain activity: A mechanistic whole-brain model of empirical MEG data. NeuroImage, 152, 538–550. 10.1016/j.neuroimage.2017.03.023

Deco, G., & Kringelbach, M. L. (2016). Metastability and Coherence: Extending the Communication through Coherence Hypothesis Using A Whole-Brain Computational Perspective. Trends in Neurosciences, 39(3), Article 3. 10.1016/j.tins.2016.01.001

Drummond, S. P. A., Brown, G. G., Gillin, J. C., Stricker, J. L., Wong, E. C., & Buxton, R. B. (2000). Altered brain response to verbal learning following sleep deprivation. Nature, 403(6770), 655–657. 10.1038/35001068

Duffy, J. F., Willson, H. J., Wang, W., & Czeisler, C. A. (2009). Healthy older adults better tolerate sleep deprivation than young adults. Journal of the American Geriatrics Society, 57(7), 1245–1251. 10.1111/j.1532-5415.2009.02303.x

Dunn, J. C. (1973). A Fuzzy Relative of the ISODATA Process and Its Use in Detecting Compact Well-Separated Clusters. Journal of Cybernetics, 3(3), 32–57. 10.1080/01969727308546046

Edinger, J. D., Glenn, D. M., Bastian, L. A., & Marsh, G. R. (2000). Slow-wave sleep and waking cognitive performance II: Findings among middle-aged adults with and without insomnia complaints. Physiology & Behavior, 70(1), 127–134. 10.1016/S0031-9384(00)00238-9

Fang, H., Tu, S., Sheng, J., & Shao, A. (2019). Depression in sleep disturbance: A review on a bidirectional relationship, mechanisms and treatment. Journal of Cellular and Molecular Medicine, 23(4), 2324–2332. 10.1111/jcmm.14170

Feinberg, I. (2000). Slow Wave Sleep and Release of Growth Hormone. JAMA, 284(21), 2717– 2718. 10.1001/jama.284.21.2717

Ferrie, J. E., Shipley, M. J., Akbaraly, T. N., Marmot, M. G., Kivimäki, M., & Singh-Manoux, A. (2011). Change in sleep duration and cognitive function: Findings from the Whitehall II Study. Sleep, 34(5), 565–573. 10.1093/sleep/34.5.565

Frazier-Logue, N., Wang, J., Wang, Z., Sodums, D., Khosla, A., Samson, A. D., McIntosh, A. R., & Shen, K. (2022). A Robust Modular Automated Neuroimaging Pipeline for Model Inputs to TheVirtualBrain. Frontiers in Neuroinformatics, 16, 883223. 10.3389/fninf.2022.883223

Gerhardsson, A., Fischer, H., Lekander, M., Kecklund, G., Axelsson, J., Åkerstedt, T., & Schwarz, J. (2019). Positivity Effect and Working Memory Performance Remains Intact in Older Adults After Sleep Deprivation. Frontiers in Psychology, 10. 10.3389/fpsyg.2019.00605

Germain, A., Buysse, D. J., & Nofzinger, E. (2008). Sleep-specific mechanisms underlying posttraumatic stress disorder: Integrative review and neurobiological hypotheses. Sleep Medicine Reviews, 12(3), 185–195. 10.1016/j.smrv.2007.09.003

Glerean, E., Salmi, J., Lahnakoski, J. M., Jääskeläinen, I. P., & Sams, M. (2012). Functional Magnetic Resonance Imaging Phase Synchronization as a Measure of Dynamic Functional Connectivity. Brain Connectivity, 2(2), 91–101. 10.1089/brain.2011.0068

Griffanti, L., Salimi-Khorshidi, G., Beckmann, C. F., Auerbach, E. J., Douaud, G., Sexton, C. E., Zsoldos, E., Ebmeier, K. P., Filippini, N., Mackay, C. E., Moeller, S., Xu, J., Yacoub, E., Baselli, G., Ugurbil, K., Miller, K. L., & Smith, S. M. (2014). ICA-based artefact and accelerated fMRI acquisition for improved Resting State Network imaging. NeuroImage, 95, 232. 10.1016/j.neuroimage.2014.03.034

Heisz, J. J., Gould, M., & McIntosh, A. R. (2015). Age-related Shift in Neural Complexity Related to Task Performance and Physical Activity. Journal of Cognitive Neuroscience, 27(3), Article 3. 10.1162/jocn_a_00725

Hemmeter, U., Bischof, R., Hatzinger, M., Seifritz, E., & Holsboer-Trachsler, E. (1998). Microsleep during Partial Sleep Deprivation in Depression. Biological Psychiatry, 43(11), 829–839. 10.1016/S0006-3223(97)00297-7

Hemmeter, U.-M., Hemmeter-Spernal, J., & Krieg, J.-C. (2010). Sleep deprivation in depression. Expert Review of Neurotherapeutics, 10(7), 1101–1115. 10.1586/ern.10.83

Huang, N.-X., Gao, Z.-L., Lin, J.-H., Lin, Y.-J., & Chen, H.-J. (2022). Altered stability of brain functional architecture after sleep deprivation: A resting-state functional magnetic resonance imaging study. Frontiers in Neuroscience, 16, 998541. 10.3389/fnins.2022.998541

Hutchison, R. M., Womelsdorf, T., Allen, E. A., Bandettini, P. A., Calhoun, V. D., Corbetta, M., Penna, S. D., Duyn, J. H., Glover, G. H., Gonzalez-Castillo, J., Handwerker, D. A., Keilholz, S., Kiviniemi, V., Leopold, D. A., de Pasquale, F., Sporns, O., Walter, M., & Chang, C. (2013). Dynamic functional connectivity: Promise, issues, and interpretations. NeuroImage, 80, 10.1016/j.neuroimage.2013.05.079. 10.1016/j.neuroimage.2013.05.079

Jenkinson, M., Beckmann, C. F., Behrens, T. E. J., Woolrich, M. W., & Smith, S. M. (2012). FSL. NeuroImage, 62(2), Article 2. 10.1016/j.neuroimage.2011.09.015

Jin, H., Ranasinghe, K. G., Prabhu, P., Dale, C., Gao, Y., Kudo, K., Vossel, K., Raj, A., Nagarajan, S. S., & Jiang, F. (2023). Dynamic functional connectivity MEG features of Alzheimer’s disease. NeuroImage, 281, 120358. 10.1016/j.neuroimage.2023.120358

Kovacevic, N., Abdi, H., Beaton, D., & McIntosh, A. R. (2013). Revisiting PLS Resampling: Comparing Significance Versus Reliability Across Range of Simulations. In H. Abdi, W. W. Chin, V. Esposito Vinzi, G. Russolillo, & L. Trinchera (Eds.), New Perspectives in Partial Least Squares and Related Methods (pp. 159–170). Springer. 10.1007/978-1-4614-8283-3_10

Krakow, B., Artar, A., Warner, T. D., Melendrez, D., Johnston, L., Hollifield, M., Germain, A., & Koss, M. (2000). Sleep disorder, depression, and suicidality in female sexual assault survivors. Crisis, 21(4), 163–170. 10.1027//0227-5910.21.4.163

Krieg, E. F., Chrislip, D. W., Letz, R. E., Otto, D. A., Crespo, C. J., Stephen Brightwell, W., & Ehrenberg, R. L. (2001). Neurobehavioral test performance in the third National Health and Nutrition Examination Survey. Neurotoxicology and Teratology, 23(6), 569–589. 10.1016/S0892-0362(01)00177-5

Kronholm, E. (2012). Sleep in cognitive life-time trajectory. Sleep Medicine, 13(7), 777–778. 10.1016/j.sleep.2012.04.001

Kulmala, J., von Bonsdorff, M. B., Stenholm, S., Törmäkangas, T., von Bonsdorff, M. E., Nygård, C.-H., Klockars, M., Seitsamo, J., Ilmarinen, J., & Rantanen, T. (2013). Perceived Stress Symptoms in Midlife Predict Disability in Old Age: A 28-Year Prospective Cohort Study. The Journals of Gerontology: Series A, 68(8), 984–991. 10.1093/gerona/gls339

Littlejohns, T. J., Holliday, J., Gibson, L. M., Garratt, S., Oesingmann, N., Alfaro-Almagro, F., Bell, J. D., Boultwood, C., Collins, R., Conroy, M. C., Crabtree, N., Doherty, N., Frangi, A. F., Harvey, N. C., Leeson, P., Miller, K. L., Neubauer, S., Petersen, S. E., Sellors, J., … Allen, N. E. (2020). The UK Biobank imaging enhancement of 100,000 participants: Rationale, data collection, management and future directions. Nature Communications, 11(1), Article 1. 10.1038/s41467-020-15948-9

Liu, T. T., Nalci, A., & Falahpour, M. (2017). The global signal in fMRI: Nuisance or Information? NeuroImage, 150, 213–229. 10.1016/j.neuroimage.2017.02.036

Lobo, A., López-Antón, R., de-la-Cámara, C., Quintanilla, M. A., Campayo, A., Saz, P., & ZARADEMP Workgroup. (2008). Non-cognitive psychopathological symptoms associated with incident mild cognitive impairment and dementia, Alzheimer’s type. Neurotoxicity Research, 14(2–3), 263–272. 10.1007/BF03033815

Luo, A. C., Sydnor, V. J., Pines, A., Larsen, B., Alexander-Bloch, A. F., Cieslak, M., Covitz, S., Chen, A. A., Esper, N. B., Feczko, E., Franco, A. R., Gur, R. E., Gur, R. C., Houghton, A., Hu, F., Keller, A. S., Kiar, G., Mehta, K., Salum, G. A., … Satterthwaite, T. D. (2024). Functional connectivity development along the sensorimotor-association axis enhances the cortical hierarchy. Nature Communications, 15(1), 3511. 10.1038/s41467-024-47748-w

Lutkenhoff, E. S., Rosenberg, M., Chiang, J., Zhang, K., Pickard, J. D., Owen, A. M., & Monti, M. M. (2014). Optimized brain extraction for pathological brains (optiBET). PloS One, 9(12), Article 12. 10.1371/journal.pone.0115551

Markello, R. D., Hansen, J. Y., Liu, Z.-Q., Bazinet, V., Shafiei, G., Suárez, L. E., Blostein, N., Seidlitz, J., Baillet, S., Satterthwaite, T. D., Chakravarty, M. M., Raznahan, A., & Misic, B. (2022). neuromaps: Structural and functional interpretation of brain maps. Nature Methods, 19(11), 1472–1479. 10.1038/s41592-022-01625-w

McCrae, C. S., Vatthauer, K. E., Dzierzewski, J. M., & Marsiske, M. (2012). Habitual Sleep, Reasoning, and Processing Speed in Older Adults with Sleep Complaints. Cognitive Therapy and Research, 36(2), 156–164. 10.1007/s10608-011-9425-4

McIntosh, A. R., & Lobaugh, N. J. (2004). Partial least squares analysis of neuroimaging data: Applications and advances. NeuroImage, 23 *Suppl 1*, S250–263. 10.1016/j.neuroimage.2004.07.020

Medic, G., Wille, M., & Hemels, M. E. (2017). Short- and long-term health consequences of sleep disruption. Nature and Science of Sleep, 9, 151–161. 10.2147/NSS.S134864

Nemeth, D., Janacsek, K., Király, K., Londe, Z., Németh, K., Fazekas, K., Adam, I., & Csányi, A. (2013). Probabilistic sequence learning in mild cognitive impairment. Frontiers in Human Neuroscience, 7. 10.3389/fnhum.2013.00318

Neudorf, J., Shen, K., & McIntosh, A. R. (2024). Reorganization of structural connectivity in the brain supports preservation of cognitive ability in healthy aging. Network Neuroscience, 8(3), 837–859. 10.1162/netn_a_00377

Neudorf, J., Shen, K., & McIntosh, A. R. (2025). Dynamic network features of functional and structural brain networks support visual working memory in aging adults. Imaging Neuroscience, 3, IMAG.a.5. 10.1162/IMAG.a.5

Nilsonne, G., Tamm, S., d’Onofrio, P., Thuné, H. Å., Schwarz, J., Lavebratt, C., Liu, J. J., Månsson, K. N. T., Sundelin, T., Axelsson, J., Lamm, C., Petrovic, P., Fransson, P., Kecklund, G., Fischer, H., Lekander, M., & Åkerstedt, T. (2016). A multimodal brain imaging dataset on sleep deprivation in young and old humans. http://openarchive.ki.se/xmlui/handle/10616/45181

Nilsonne, G., Tamm, S., Schwarz, J., Almeida, R., Fischer, H., Kecklund, G., Lekander, M., Fransson, P., & Åkerstedt, T. (2017). Intrinsic brain connectivity after partial sleep deprivation in young and older adults: Results from the Stockholm Sleepy Brain study. Scientific Reports, 7(1), 9422. 10.1038/s41598-017-09744-7

Pace-Schott, E. F., & Spencer, R. M. C. (2011). Age-related changes in the cognitive function of sleep. Progress in Brain Research, 191, 75–89. 10.1016/B978-0-444-53752-2.00012-6

Park, D. C., & Reuter-Lorenz, P. (2009). The adaptive brain: Aging and neurocognitive scaffolding. Annual Review of Psychology, 60, 173–196. 10.1146/annurev.psych.59.103006.093656

Peters, K. R., Ray, L., Smith, V., & Smith, C. (2008). Changes in the density of stage 2 sleep spindles following motor learning in young and older adults. Journal of Sleep Research, 17(1), 23–33. 10.1111/j.1365-2869.2008.00634.x

Pilcher, J. J., & Huffcutt, A. I. (1996). Effects of Sleep Deprivation on Performance: A Meta-Analysis. Sleep, 19(4), 318–326. 10.1093/sleep/19.4.318

Ponce-Alvarez, A., Deco, G., Hagmann, P., Romani, G. L., Mantini, D., & Corbetta, M. (2015). Resting-State Temporal Synchronization Networks Emerge from Connectivity Topology and Heterogeneity. PLOS Computational Biology, 11(2), e1004100. 10.1371/journal.pcbi.1004100

Regestein, Q. R., Friebely, J., Shifren, J. L., Scharf, M. B., Wiita, B., Carver, J., & Schiff, I. (2004). Self-reported sleep in postmenopausal women. *Menopause (New York*, N.Y.), 11(2), 198–207. 10.1097/01.gme.0000097741.18446.3e

Rousseeuw, P. J. (1987). Silhouettes: A graphical aid to the interpretation and validation of cluster analysis. Journal of Computational and Applied Mathematics, 20, 53–65. 10.1016/0377-0427(87)90125-7

Rubinov, M., & Sporns, O. (2010). Complex network measures of brain connectivity: Uses and interpretations. NeuroImage, 52(3), Article 3. 10.1016/j.neuroimage.2009.10.003

Salas, R. E., Galea, J. M., Gamaldo, A. A., Gamaldo, C. E., Allen, R. P., Smith, M. T., Cantarero, G., Lam, B. D., & Celnik, P. A. (2014). Increased Use-Dependent Plasticity in Chronic Insomnia. Sleep, 37(3), 535–544. 10.5665/sleep.3492

Schaefer, A., Kong, R., Gordon, E. M., Laumann, T. O., Zuo, X.-N., Holmes, A. J., Eickhoff, S. B., & Yeo, B. T. T. (2018). Local-Global Parcellation of the Human Cerebral Cortex from Intrinsic Functional Connectivity MRI. Cerebral Cortex (New York, N.Y.: 1991), 28(9), Article 9. 10.1093/cercor/bhx179

Schmidt, C., Peigneux, P., & Cajochen, C. (2012). Age-related changes in sleep and circadian rhythms: Impact on cognitive performance and underlying neuroanatomical networks. Frontiers in Neurology, 3, 118. 10.3389/fneur.2012.00118

Schwarz, J., Axelsson, J., Gerhardsson, A., Tamm, S., Fischer, H., Kecklund, G., & Åkerstedt, T. (2019). Mood impairment is stronger in young than in older adults after sleep deprivation. Journal of Sleep Research, 28(4), e12801. 10.1111/jsr.12801

Scullin, M. K. (2013). Sleep, Memory, and Aging: The Link Between Slow-Wave Sleep and Episodic Memory Changes from Younger to Older Adults. Psychology and Aging, 28(1), 105–114. 10.1037/a0028830

Scullin, M. K., & Bliwise, D. L. (2015). Sleep, Cognition, and Normal Aging: Integrating a Half-Century of Multidisciplinary Research. Perspectives on Psychological Science : A Journal of the Association for Psychological Science, 10(1), 97. 10.1177/1745691614556680

Shehzad, Z., Kelly, C., Reiss, P. T., Cameron Craddock, R., Emerson, J. W., McMahon, K., Copland, D. A., Xavier Castellanos, F., & Milham, M. P. (2014). A multivariate distance-based analytic framework for connectome-wide association studies. NeuroImage, 93, 74–94. 10.1016/j.neuroimage.2014.02.024

Song, J., Birn, R. M., Boly, M., Meier, T. B., Nair, V. A., Meyerand, M. E., & Prabhakaran, V. (2014). Age-Related Reorganizational Changes in Modularity and Functional Connectivity of Human Brain Networks. Brain Connectivity, 4(9), 662–676. 10.1089/brain.2014.0286

Spencer, R. M. C., & Pace-Schott, E. F. (2013). Age-related changes in consolidation of perceptual and muscle-based learning of motor skills. Frontiers in Aging Neuroscience, 5. 10.3389/fnagi.2013.00083

Spiegel, R., Koberle, S., & Allen, S. R. (1986). Significance of Slow Wave Sleep: Considerations from a Clinical Viewpoint. Sleep, 9(1), 66–79. 10.1093/sleep/9.1.66

Spira, A. P., Gamaldo, A. A., An, Y., Wu, M. N., Simonsick, E. M., Bilgel, M., Zhou, Y., Wong, D. F., Ferrucci, L., & Resnick, S. M. (2013). Self-reported sleep and β-amyloid deposition in community-dwelling older adults. JAMA Neurology, 70(12), 1537–1543. 10.1001/jamaneurol.2013.4258

Sporns, O. (2018). Graph theory methods: Applications in brain networks. Dialogues in Clinical Neuroscience, 20(2), Article 2.

Stenfors, C. U. D., Hanson, L. M., Oxenstierna, G., Theorell, T., & Nilsson, L.-G. (2013). Psychosocial Working Conditions and Cognitive Complaints among Swedish Employees. PLOS ONE, 8(4), e60637. 10.1371/journal.pone.0060637

Sternberg, D. A., Ballard, K., Hardy, J. L., Katz, B., Doraiswamy, P. M., & Scanlon, M. (2013). The largest human cognitive performance dataset reveals insights into the effects of lifestyle factors and aging. Frontiers in Human Neuroscience, 7. 10.3389/fnhum.2013.00292

Sutter, C., Zöllig, J., Allemand, M., & Martin, M. (2012). Sleep quality and cognitive function in healthy old age: The moderating role of subclinical depression. Neuropsychology, 26(6), 768–775. 10.1037/a0030033

Tian, Y., Margulies, D. S., Breakspear, M., & Zalesky, A. (2020). Topographic organization of the human subcortex unveiled with functional connectivity gradients. Nature Neuroscience, 23(11), Article 11. 10.1038/s41593-020-00711-6

Tian, Y., Peng, X.-R., Tang, Z., Long, Z., Xie, C., & Lei, X. (2024). Enhanced diversity on connector hubs following sleep deprivation: Evidence from diffusion and functional magnetic resonance imaging. NeuroImage, 299, 120837. 10.1016/j.neuroimage.2024.120837

Virta, J. J., Heikkilä, K., Perola, M., Koskenvuo, M., Räihä, I., Rinne, J. O., & Kaprio, J. (2013). Midlife sleep characteristics associated with late life cognitive function. Sleep, 36(10), 1533–1541, 1541A. 10.5665/sleep.3052

Wu, J., Ngo, G. H., Greve, D., Li, J., He, T., Fischl, B., Eickhoff, S. B., & Yeo, B. T. T. (2018). Accurate nonlinear mapping between MNI volumetric and FreeSurfer surface coordinate systems. Human Brain Mapping, 39(9), 3793–3808. 10.1002/hbm.24213

Xu, H., Shen, H., Wang, L., Zhong, Q., Lei, Y., Yang, L., Zeng, L.-L., Zhou, Z., Hu, D., & Yang, Z. (2018). Impact of 36 h of total sleep deprivation on resting-state dynamic functional connectivity. Brain Research, 1688, 22–32. 10.1016/j.brainres.2017.11.011

Yeo, B. T. T., Tandi, J., & Chee, M. W. L. (2015). Functional connectivity during rested wakefulness predicts vulnerability to sleep deprivation. NeuroImage, 111, 147–158. 10.1016/j.neuroimage.2015.02.018

Zalesky, A., Fornito, A., & Bullmore, E. T. (2010). Network-based statistic: Identifying differences in brain networks. NeuroImage, 53(4), Article 4. 10.1016/j.neuroimage.2010.06.041

Zang, F., Liu, X., Fan, D., He, C., Zhang, Z., Xie, C., Initiative, for the A. D. N., & Consortium, for the A. D. M. (2024). Dynamic functional network connectivity and its association with lipid metabolism in Alzheimer’s disease. CNS Neuroscience & Therapeutics, 30(9), e70029. 10.1111/cns.70029

Zhou, X., Wu, T., Yu, J., & Lei, X. (2017). Sleep Deprivation Makes the Young Brain Resemble the Elderly Brain: A Large-Scale Brain Networks Study. Brain Connectivity, 7(1), 58–68. 10.1089/brain.2016.0452

